# On the prediction of human intelligence from neuroimaging: A systematic review of methods and reporting

**DOI:** 10.1101/2021.10.19.462649

**Authors:** Bruno Hebling Vieira, Gustavo Santo Pedro Pamplona, Karim Fachinello, Alice Kamensek Silva, Maria Paula Foss, Carlos Ernesto Garrido Salmon

## Abstract

Human intelligence is one of the main objects of study in cognitive neuroscience. Reviews and meta-analyses have proved to be fundamental to establish and cement neuroscientific theories on intelligence. The prediction of intelligence using in vivo neuroimaging data and machine learning has become a widely accepted and replicated result. Here, we present a systematic review of this growing area of research, based on studies that employ structural, functional, and/or diffusion MRI to predict human intelligence in cognitively normal subjects using machine-learning. We performed a systematic assessment of methodological and reporting quality, using the PROBAST and TRIPOD assessment forms and 30 studies identified through a systematic search. We observed that fMRI is the most employed modality, resting-state functional connectivity (RSFC) is the most studied predictor, and the Human Connectome Project is the most employed dataset. A meta-analysis revealed a significant difference between the performance obtained in the prediction of general and fluid intelligence from fMRI data, confirming that the quality of measurement moderates this association. The expected performance of studies predicting general intelligence from fMRI was estimated to be *r* = 0.42 (CI_95%_ = [0.35, 0.50]) while for studies predicting fluid intelligence obtained from a single test, expected performance was estimated as *r* = 0.15 (CI_95%_ = [0.13, 0.17]). We further enumerate some virtues and pitfalls we identified in the methods for the assessment of intelligence and machine learning. The lack of treatment of confounder variables, including kinship, and small sample sizes were two common occurrences in the literature which increased risk of bias. Reporting quality was fair across studies, although reporting of results and discussion could be vastly improved. We conclude that the current literature on the prediction of intelligence from neuroimaging data is reaching maturity. Performance has been reliably demonstrated, although extending findings to new populations is imperative. Current results could be used by future works to foment new theories on the biological basis of intelligence differences.

## 1. Introduction

Intelligence is a broad construct comprising multiple components, which can be estimated with a range of well-established tests (Urbina 2011). Regardless of the instrument, scores in intelligence tests are positively correlated. G was postulated to be the “general factor” explaining this phenomenon by Spearman (1904), whose evidence “[…] can be said to be overwhelming” (Carroll 1997). Albeit originally terming it “general intelligence”, Spearman later in his life adopted a critical view of the term and ceased to associate it with G (Spearman 1927). Henceforth, to avoid ambiguities in this review we will employ the widely used term “intelligence”. Even though G successfully captures the overall positive correlation (Spearman 1904), there is controversy regarding its validity as a single, all-encompassing, measure of intelligence. An alternative view posits that intelligence comprises multiple factors (Thurstone 1938). Posteriorly, an integrated model for intelligence called **G^F^**-**G^C^** was proposed by Cattell (1941, 1971). **G^F^** stands for fluid intelligence and is associated with inductive and deductive reasoning, covering non-verbal components; therefore, it does not depend on previously acquired knowledge and the influence of culture. Concept formation and recognition, identification of complex relationships, understanding of implications, and making inferences are examples of tasks related to **G^F^**. On the other hand, **G^C^**, crystallized intelligence, comprises the knowledge acquired through life experience and education related to cultural experiences. Hence, crystallized capacities are demonstrated, for example, in tasks regarding the recognition of the meaning of words (Schelini 2006). While the scientific construction of G is based on correlations between test scores, intelligence quotient (IQ) is based on the sum of standardized scores of commonly used cognitive batteries, such as Wechsler scales with full scale IQ (FSIQ), verbal IQ (VIQ), and performance IQ (PIQ). FSIQ scores are excellent measures of G (Gignac et al. 2017) representing the general level of cognitive functioning. VIQ relates to verbal comprehension, acquired knowledge, language processing, verbal reasoning, attention, verbal learning, and memory. In sharp contrast, PIQ is connected to perceptual organization, processing visual, planning ability, non-learning-verbal and thinking skills, and manipulating visual stimuli with speed.

Studies show associations between brain and behavior measurements. The first finding was the positive correlation between brain volume or intracranial volume and intelligence (Luders et al. 2009; McDaniel 2005). Other structural MRI (sMRI) correlates of intelligence include fine-grained morphometry, such as callosal thickness (Luders et al. 2007), striatal volume (Grazio-plene et al. 2015) and regional gray and white matter volumetry (Haier et al. 2005). Functional connectivity (FC), as measured by functional MRI (fMRI), has reliably been shown to correlate with G and IQ. This includes correlations between resting-state FC (RSFC) network organization and FSIQ (Pamplona et al. 2015; Song et al. 2008) and regional global connectivity and **G^F^** (Cole et al. 2012). The topography of task fMRI (T-fMRI) statistical maps have been found to correlate with intelligence as well (Choi et al. 2008; Graham et al. 2010). Correlates of intelligence extend beyond fMRI RSFC and task activations as well, to include measures such as amplitude of low frequency fluctuations (ALFF) and dynamic functional connectivity (dynFC). Using multimodal magnetic resonance imaging (MRI) Ritchie et al. (2015) demonstrates a plethora of correlates of G, including diffusion MRI (dMRI). For extensive literature reviews, see Basten et al. (2021) and Dizaji et al. (2021).

Previous reviews (Barbey 2018; Jung et al. 2007) and meta-analyses (Basten et al. 2015; McDaniel 2005; Pietschnig et al. 2015) were fundamental in the development of theories of biological intelligence. At the time studies performing predictive analyses were scarcer than today. This type of analysis enjoys growing popularity in neuroimaging (Bzdok 2017; Bzdok et al. 2018). machine learning (ML)-based predictive analyses allow one to test a much more complex hypothesis space than univariate, group-based testing. The multivariate nature of ML allows interactions and commonalities between predictors to be taken into account. It also “tests” such hypotheses on the basis of individualized predictions, taking into account heterogeneity that is diluted in group-based analyses (Sui et al. 2020). Data-driven studies based on ML are fundamental to understand the degree that variability in brain phenotypes explain variability in intelligence. ML-based studies also address the question of generalizability patterns at the forefront. For these reasons, this type of study is widely used in the investigation of behavior, with cognition and, specifically, human intelligence as the most studied domains (Sui et al. 2020).

While the literature of brain correlates on intelligence covers various techniques, such as sMRI, fMRI, dMRI, positron emission tomography (PET), electroencephalography (EEG), magnetoencephalography (MEG), predictive studies are limited in this regard. Availability is one of the main factors behind that choice, because ML benefits from large amounts of data (Cui et al. 2018). Small data has been identified as one source of optimistic bias in error-bars (Varoquaux 2018), and leads to non-reproducible results. Large-scale open-data imaging cohorts are often centered on fMRI, with sMRI and dMRI providing complimentary information. For this reason, we opted to focus on fMRI, sMRI and dMRI, anticipating a small incidence of studies using other imaging modalities.

A large number of studies on the prediction of intelligence was published in recent years. To the best of the authors’ knowledge, no review on this application of ML to predict human intelligence from brain imaging has been previously published. The purpose of this review is to identify existing literature, critically appraise reporting and methodology. We hope that our work will promote the establishment of best practices and prospects for future research in this field of research.

## 2. Methods

This review was developed following Preferred Reporting Items for Systematic Reviews and Meta-Analyses (PRISMA) guidelines for transparent reporting of systematic reviews (Moher et al. 2009). See Table B.4 for the PRISMA checklist. Choice of methods and search strategy are based on a protocol we developed and registered at Open Science Framework (Vieira et al. 2021a). Post-hoc adaptations are mentioned below, when applicable.

### 2.1. Eligibility criteria

Eligibility criteria were peer-reviewed original articles written in English that performed individualized prediction of intelligence using at least one of fMRI, sMRI and dMRI in neurotypical human subjects using ML and include evaluation of generalizability, *i.e.* cross-validation, bootstrapping, or external validation.

### 2.2. Information sources

We performed a systematized search in Scopus (scopus.com), dating to 15th December 2020. Additional documents were retrieved from a recent literature review (Dizaji et al. 2021), co-authored by B.H.V. and C.E.G.S., and another study (Fan et al. 2020, Table 1) that provide a comparison between similar studies.

**Table 1:** General characteristics of documents retrieved using based on our data extraction form. “a”: primarily about predictive modeling; “b”: primarily about intelligence;

| Studies | Number of subjects | Input | ML models | Validation strategy | Target |
| --- | --- | --- | --- | --- | --- |
| Choi et al. 2008<sup>b</sup> | 408 for FA, 225 for prediction from NRI/KAIST (training data: 116 sMRIs and 61 fMRIs; test data: 48); | fMRI & sMRI<br>Cortical thickness, T-fMRI activation in a fluid reasoning task, gray matter volume, sex | Linear modelling (derived from separate structural and functional samples) | Independent test sample | G |
| Yang et al. 2013<sup>ab</sup> | 78 from NRI | sMRI<br>Cortical thickness, surface area, sulcal depth, mean curvature | PLSR | LOOCV | FSIQ |
| Finn et al. 2015 | 118 from HCP (Q2 release) | fMRI<br>RSFC under various preprocessing pipelines | CPM | LOOCV | G<sub>F</sub> |
| Wang et al. 2015<sup>ab</sup> | 164 from ABIDE | sMRI<br>Regional gray and white matter volume | Multi-kernel KSVR following multiple feature selection | Repeated (10x) 10-fold CV, with inner CV (unspecified) for parameter tuning | IQ |
| Ferguson et al. 2017<sup>b</sup> | 830 from HCP (S900 release) (600 for training, 230 for testing) | fMRI<br>Scaled eigenvalues from spectral decomposition of concatenated RS-fMRI, products of eigenvalues | LASSO | Independent test sample | G<sub>F</sub> |
| Powell et al. 2017<sup>a</sup> | 841 from HCP | dMRI<br>Local Connectome Fingerprints and intracranial volume | LASSO PCR | 5-fold CV | G<sub>F</sub> |
| Greene et al. 2018<sup>a</sup> | 515 from HCP; 571 from PNC | fMRI<br>RSFC and T-fMRI activation (7 tasks in HCP, 2 in PNC) | CPM | LOOCV (within samples) and between samples/between conditions validation | G<sub>F</sub> |
| Dubois et al. 2018<sup>b</sup><sup>a</sup> | 884 from HCP (S1200 release) | fMRI<br>RSFC under various preprocessing pipelines | CPM & elastic net following univariate filtering | LOFOCV (410 families) | G<sub>F</sub> |
| Dubois et al. 2018<sup>a</sup><sup>ab</sup> | 884 for CV, 1181 for FA from HCP | fMRI<br>RSFC | Elastic Net after univariate filtering | LOFOCV | G |
| Li et al. 2018<sup>ab</sup> | 100 from HCP (Unrelated subjects) | fMRI<br>ALFF following voxelwise univariate filtering, seed-based FC | L2SVR | LOOCV | G<sub>F</sub> |
| Cox et al. 2019<sup>b</sup> | 27100 for FA and 4768 for training, 2510 for testing with fractional anisotropy; 4707 for training, 2494 for testing with mean diffusion; cortical: 5246 for training, 2589 for testing with cortical volume; 5253 for training, 2595 for testing with subcortical volume; from the UK Biobank | sMRI & dMRI<br>ROI white matter mean diffusivity and fractional anisotropy and gray matter cortical and subcortical volumes | MIMIC | Independent test sample (Manchester = training data, Newcastle = test data) | G |
| Yang et al. 2019<sup>b</sup> | 68 from HCP-Q1 | fMRI<br>RS-fMRI temporal variances of temporal autocorrelations (sulci, gyri, undefined cortices) from four ROIs | Linear regression | LOOCV | G<sub>F</sub> |

Table 1 continued from previous page
| Studies | Number of subjects | Input | ML models | Validation strategy | Target |
| --- | --- | --- | --- | --- | --- |
| Zhang et al. 2019 | 1065 from HCP | dMRI & fMRI<br>Structural connectivity tensor (weighted according to 12 factors based on diffusion, endpoints and geometry); RSFC; local structural connectivity | Linear regression<br>(after tensor network PCA with k = 60) | 5-fold CV | G<sub>F</sub> |
| Gao et al. 2019<sup>a</sup> | 515 from HCP; 571 from PNC | fMRI<br>RSFC and T-fMRI FC | rCPM, GFC-ridge,<br>cCPM, CPM, GFC-CPM | Repeated (100x) 10-fold CV; External Validation | G<sub>F</sub> |
| Dadi et al. 2019<sup>a</sup> | 443 from HCP (213 High IQ, 230 Low IQ, based on terciles) | fMRI<br>RSFC | K-Nearest Neighbors (K = 1, Euclidean distance metric), Gaussian Naïve Bayes, Random Forests, L1-SVC L1-LogReg, Ridge classification, L2SVC, L2LogReg, 10%-univariate ANOVA SVC | Repeated (100x) Stratified Holdout (75%) | G<sub>F</sub> |
| Elliott et al. 2019 | 298 from HCP; 591 from Dunedin Study | fMRI<br>RSFC, GFC | CPM | LOOCV (within samples) and between samples validation | Cognitive Ability |
| Yoo et al. 2019 | 316 unrelated subjects (out of 563) from HCP (S1200 release) | fMRI<br>Bivariate and multivariate (distance correlation) RSFC | CPM | Repeated (5000x) 10-fold CV | G<sub>F</sub> |
| Li et al. 2019 | 862 from BGSP, 953 from HCP | fMRI<br>RSFC | KRR (correlation kernel) | 20-fold nested family-aware CV, inner 20-fold CV for selection of parameters | G<sub>F</sub> |
| Kashyap et al. 2019<sup>a</sup> | 803 from HCP | fMRI<br>RSFC with and without Common Orthogonal Basis Extraction (COBE) | Elastic Net after univariate filtering | 20-fold nested CV, with inner CV for tuning | G<sub>F</sub> |
| Kong et al. 2019<sup>a</sup> | 577 from HCP | fMRI<br>Dice overlap kernel of different parcellation algorithms (ICA back-projection algorithm, individual-specific parcellation algorithm of Gordon, parcellation algorithm of Wang, multi-session hierarchical Bayesian model (MS-HBM)) | KRR (dice overlap kernel) | Repeated (100x) 20-fold family-aware CV nested with inner tuning 20-fold CV | G<sub>F</sub> |
| Dryburgh et al. 2020<sup>ab</sup> | 226 from ABIDE-I | fMRI<br>RSFC | CPM | LOOCV | FSIQ and Verbal intelligence quotient (VIQ) |
| Jiang et al. 2020<sup>b</sup> | 326 from UESTC | fMRI & sMRI<br>RSFC, cortical thickness (vertexwise) | CPM | LOOCV | FSIQ |
| Hilger et al. 2020<sup>ab</sup> | 308 from NKI (Enhanced) | sMRI<br>Gray matter volume (voxel and regionwise) | PCA-SVR & Atlas-SVR | 10-fold CV (with nested inner 3-fold CV for parameter tuning) stratified for intelligence | FSIQ |
| Fan et al. 2020<sup>ab</sup> | 1050 from HCP | fMRI<br>dynFC | Deep neural network (CNN-LSTM), SVR | 10-fold CV (no splitting runs from the same subject) | G<sub>F</sub> and Crystallized Intelligence |

| Studies | Number of subjects | Input | ML models | Validation strategy | Target |
| --- | --- | --- | --- | --- | --- |
| Sripada et al. 2020<sup>ab</sup> | 967 for **T-fMRI**, 903 for **RS-fMRI** from **HCP** (S1200 release) | Brain Basis Set (BBS) modeling decomposition of task contrasts, **fMRI** **RSFC** | Linear regression (75 components/coefficients) | 10-fold family-aware **CV** | General Cognitive Ability (computed for each fold) |
| Wei et al. 2020<sup>a</sup> | 1003 (812 “recon2” used as discovery set; 191 “recon1” as validation set) from **HCP** (S1200 release) | **fMRI** **RSFC** | **CPM**, **SVR**, **LASSO**, and Ridge regression, after Bootstrapping Feature selection | 10-fold stratified **CV** & independent validation set | **G<sub>F</sub>** |
| He et al. 2020<sup>a</sup> | 953 from **HCP** (S1200 release); 8868 from UK Biobank | **fMRI** **RSFC** | **KRR**, **FNN**, Brain-NetCNN, **GNN** | **HCP**: 20-fold family-aware **CV** nested with inner tuning; UK Biobank: Holdout (6868 training, 1000 validation and 1000 test) | **G<sub>F</sub>** |
| Jiang et al. 2020a<sup>ab</sup> | 360 from **UESTC**; 200 from **HCP** (Q3 release); 120 from **COBRE** (60 **HCS**) | **fMRI** **RSFC** | **LASSO** | **LOOCV** (with nested 10-fold **CV** for tuning) | **FSIQ** and **G<sub>F</sub>** |
| Wu et al. 2020 | 922 from **HCP** (S1200 release) (830 for training and 92 for test) | **T-fMRI** activation in seven **HCP** tasks (emotion, gambling, language, motor, relational, social, and working memory) **fMRI** | **PLSR** | Independent test sample | **G<sub>F</sub>** and Fluid, Crystallized and Total Scores |
| Lin et al. 2020<sup>a</sup> | 143 (1 subject with missing data) from **HCP** (S900 release) | **dMRI** & **fMRI** **RSFC** and structural connectivity (quantitative anisotropy, mean streamline length, and normalized number of streamlines) | **CPM** | **LOOCV** | **G<sub>F</sub>** |

### 2.3. Search strategy

We retrieved all documents in Scopus that contained at least one of the following terms in their title, abstract, or keywords: “morphometry”, “cortical thickness”, “functional connectivity”, “structural connectivity”, or “effective connectivity”. Simultaneously, the document should contain at least one of the following terms: “prediction”, “predict”, “CPM”, “multivariate pattern analysis”, “bases”, “variability”, or “mvpa”. The documents should also contain in their title one of the following terms: “intelligence”, “behavioral”, “behavior”, “cognitive ability” or “cognition”. See Appendix A for the actual search string used.

After removal of duplicates, all records had title and abstract screened. Records were discarded if we could identify disagreement with inclusion criteria, and kept otherwise. Remaining records were retained for full-text inspection. If in accordance with the inclusion criteria, these were retained as eligible for qualitative synthesis. Otherwise discarded with reasons, *e.g.*, non-human subjects, no validation or other generalizability evaluation, did not predict intelligence, did not use neurotypical subjects.

### 2.4. Data collection process

We originally planned to use the CHecklist for critical Appraisal and data extraction for systematic Reviews of prediction Modelling Studies (CHARMS) checklist (Moons et al. 2014), but ultimately it became clear that we needed a form tailored for our research question. We constructed our own data extraction form, borrowing from CHARMS.

An online document was created and shared between authors B.H.V., K.F. and A.K.S. All three authors performed data extraction, including: (1) identification (title, year, source title, digital object identifier), (2) study population (dataset, number of subjects for psychometric assessment, number of subjects for prediction, age and sex characteristics of the sample), (3) methods (imaging modality, input features, number of features, ML models, validation strategy, performance metrics, a priori feature selection, construct, instrument, components of intelligence, scores, quality of cognitive assessment), (4) results (performance of individual methods/data combinations, best performance).

Regarding the quality of intelligence measurement, retrieved items included, when applicable: number of subtests, number of dimensions, time duration of test application, test completeness. These are important to assess whether the test properly measures intelligence and is applicable to the construct.

We additionally retrieved citations among identified documents, to build a citation network. Due to differences in formats and unreliability of automatic searching, we opted to perform a manual search over all documents. For each identified studied, we searched for the names of first authors of every other document. For consistency, we opted to consider citations of pre-print versions (same authors and title) of identified documents.

We used the Prediction model risk of bias assessment tool (PROBAST) to assess risk of bias (RoB) and concerns regarding applicability in individual studies. This was a choice made post-hoc to the registration of the study. We originally planned to create an RoB assessment checklist for the reviewed studies, but after registration we became aware of PROBAST, which fulfilled this role, requiring minimal adaptations. This assessment was performed at the result-level.

It is critical to ensure that reporting is transparent in order to ensure that findings can be replicated. We also used the Transparent Reporting of a Multivariable Prediction Model for Individual Prognosis or Diagnosis (TRIPOD) (Collins et al. 2015; Moons et al. 2015) checklist assessment tool (Heus et al. 2019) to evaluate reporting quality. We used a modified version tailored for ML predictions (Wang et al. 2020), including three modified items, shown in Table D.5. Several items in TRIPOD were not adequate for our research question, and were removed from the questionnaire for our evaluation. A few items and subitems were deemed not applicable or not important to our review question, and their assessments do not appear in this review. Namely, 1.i, 1.iii, 2.iii, 2.iv, 2.xi, 3b, 4a, 4b, 5c, 6b, 7a.iv, 7b, 10a, 10b.iv, 10b.v, 10c, 10d.ii, 10e, 11, 13a, 13b.iii, 13b.iv, 13c.ii, 15a.ii, 15b, 16.iii, 17, and 20.i, 22.ii. Items 1 and 16.i were edited to allow NA entries, due to studies that had broader scopes than the one pertaining to this review’s question. Item 13b, pertaining to demographics, requires description of the actual data being used, and not from the original sample before exclusions. Items 13b and 14a, that should be assessed based on “Results” sections, were extended to “Methods” sections as well. We performed the TRIPOD assessment at the study-level and performed across-studies summarization of reporting quality ratings.

Authors G.S.P.P. and B.H.V. completed PROBAST and TRIPOD independently. To ensure both reviewers’ interpretations were aligned, calibration was performed twice, using one study from each checklist on each occasion. Interrater agreement was then computed based on the Kappa statistic, at the score-level, for the remaining documents, excluding the two used for calibration.

The quality of measurement of intelligence is linked to validity and can interfere on results of each study. For example, Gignac et al. (2017) demonstrated that the quality of measurement moderates the association between intelligence and brain volume. The guide for categorization of measurement quality by Gignac et al. (2017) proposes four quality criteria: the number of tests, the number of group-level dimensions, testing time, and correlation with G. Authors K.F. and A.K.S. performed the assessment of measurement quality based on these criteria. “Number of tests” is categorized into 1, 1-2, 2-8, and 9+ which signal “poor”, “reasonable”, “good”, and “excellent” measures of G, respectively, in the absence of any other information. Therefore, a minimum of nine tests is needed to represent an excellent G. The “number of group-level dimensions” criterion is divided into 1, 1-2, 2-3, and 3+ test dimensions, leading to the respective classifications “poor”, “reasonable”, “good”, and “excellent” measures of G, in the absence of any other information. So, an excellent measure of G is expected to present at least three group-level dimensions, *e.g.*, **G^F^**, **G^C^**, processing speed. “Testing times” of 3-9 min, 10-19 min, 20-39 min, and 40+ minutes are respectively classified as possibly “poor”, “reasonable”, “good”, and “excellent” measures of G. The last criterion, “correlation with G”, is the best indicator of measurement quality and takes precedence over the others. However, this correlation is scarcely reported. Gignac et al. (2017) recommends substituting the correlation with G with the three other criteria.

The primary measure of prediction performance evaluation was chosen to be the Pearson correlation coefficient, R-squared and mean squared error (MSE). See Appendix C for a mathematical description of different performance measures. The Pearson correlation coefficient is the most used measure in the literature. It is scale- and location-invariant, which means that high values can be obtained with arbitrarily large errors. R-squared, when properly evaluated, is a less biased measure of explained variance than the correlation coefficient squared. However, it also suffers from its own biases that will be discussed below, requiring proper care regarding the variance of the sample. Ideally, MSE or mean absolute error (MAE) should be used when comparing different models applied to the same data (Poldrack et al. 2020). Regardless of the choice of the performance measure, comparisons between modeling approaches using different data can be ambiguous, since intrinsic variation can differ between datasets.

### 2.5. Synthesis of results

To determine the level of performance expected for each modality, we estimated a mixed-effects meta-analytic model using the package “metafor” in R 4.0.5 (Viechtbauer 2010) using results that were rated with both low RoB and low concerns regarding applicability in PROBAST. The number of samples was taken to be the total number of subjects used in the estimation of performance with pooled or unpooled means. We employed the Hunter-Schmidt estimator to deal with the sampling variance, which entails a homogeneity assumption. Different datasets and measurements of intelligence were treated as fixed effects. The same procedures were used for the R-squared, except that the Hunter-Schmidt estimator was not applied, since it pertains exclusively to correlation coefficients. Residual heterogeneity, *i.e.* the variability unaccounted for by the model and covariates, was measured by the *I*^2^statistic.

Standard errors are seldom reported in the literature. Moreover, due to the nature of cross-validation (CV), where resulting models across folds are not independent, standard errors are underestimated (Varoquaux 2018).

Assessment of within-study selective reporting is unfeasible in our setting, due to the lack of pre-registrations. Due to computational resources available today, the risk of selective reporting is real, leading to overfitting of the validation set. For an in-depth exposition, see Hosseini et al. (2020).

The funnel-plot was used to qualitatively assess the risk of publication bias.

Since one of the biggest bottlenecks for ML is sample size, we compared the number of training samples used with measured performances across studies. Training set size is often not homogeneous within studies. For CV-based studies, including leave-one-family-out CV, we chose to approximate it as *N ×* (*K −* 1)*/K*, where *N* is the total amount of data available for training and *K* is the number of groupings, *i.e.* folds or families. The formula holds true for leave-one-out CV as well. For Holdout-based studies, the actual number of training data is given by the studies.

## 3. Results

Our search strategy identified 689 records in Scopus. Additionally, 17 records were identified from Dizaji et al. (2021) and 7 in Fan et al. (2020). 63 records remained after removal of duplicates and screening. These were submitted to full-text eligibility analysis. 30 records were considered eligible for qualitative synthesis. See Figure 1. The number of studies per year is shown in Figure 2. General characteristics from each document obtained with our data extraction form are reported in Table 1.

**Figure 1:**
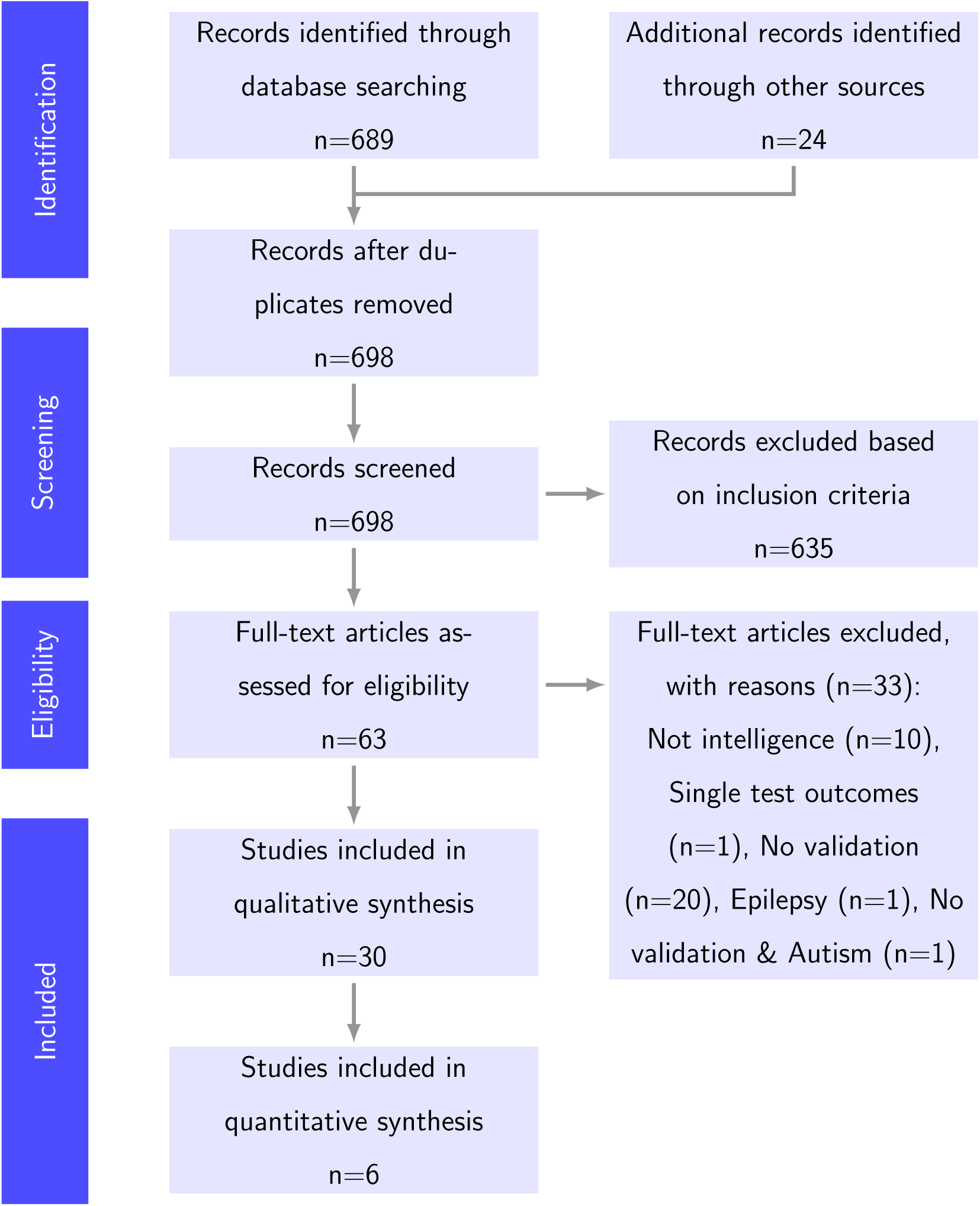
Systematic review flow diagram. See PRISMA statement (Moher et al. 2009)

**Figure 2:**
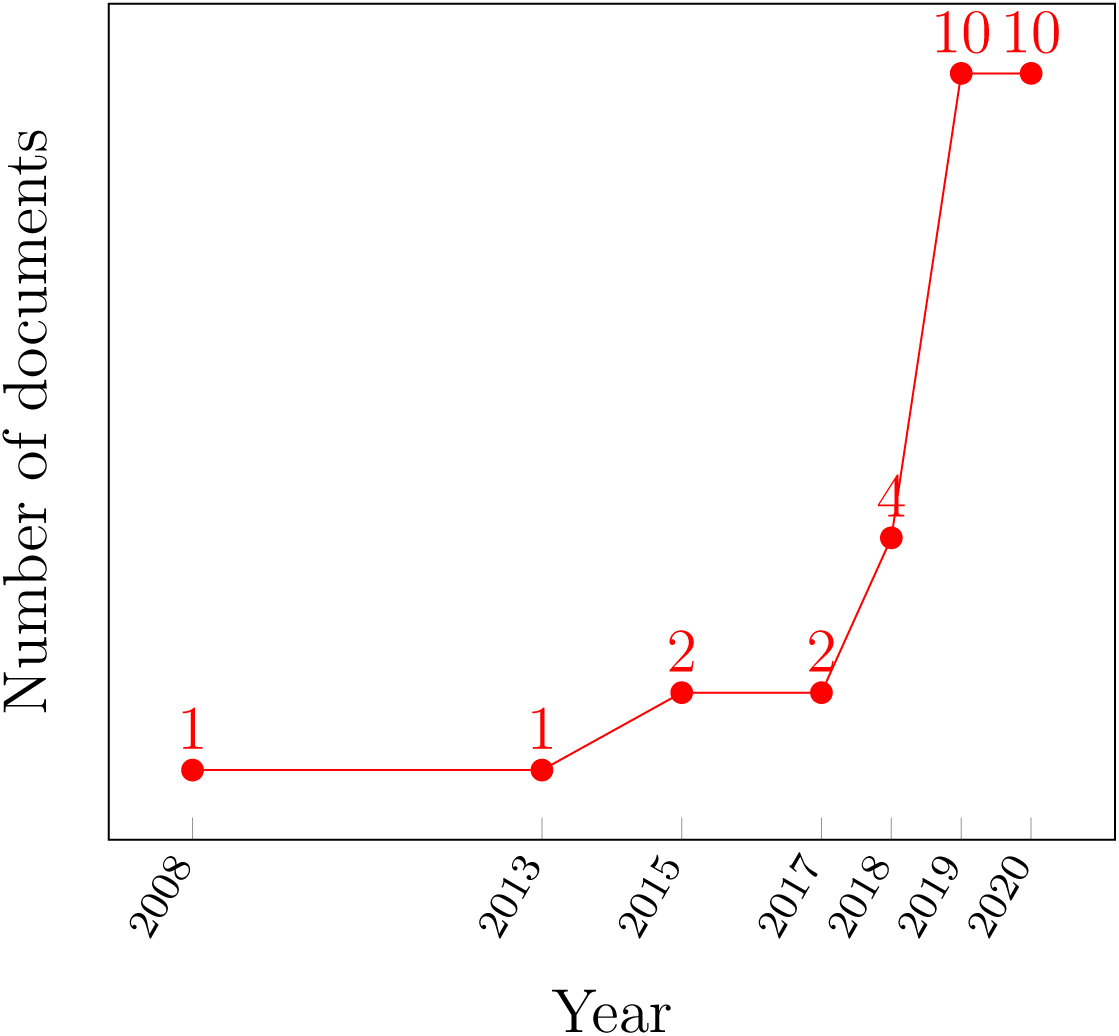
Year of publication of the 30 studies identified. An upward tendency is demonstrated, with 20 studies being published in 2019 and 2020 alone.

A co-citation network is shown in Figure 3. Arrows point from the cited to the citing document. In total, 78 citations were identified. This network systematically demarks highly influential works in the sample. Finn et al. (2015) is cited by 23 studies, out of 26 studies that were published posteriorly to it.

**Figure 3:**
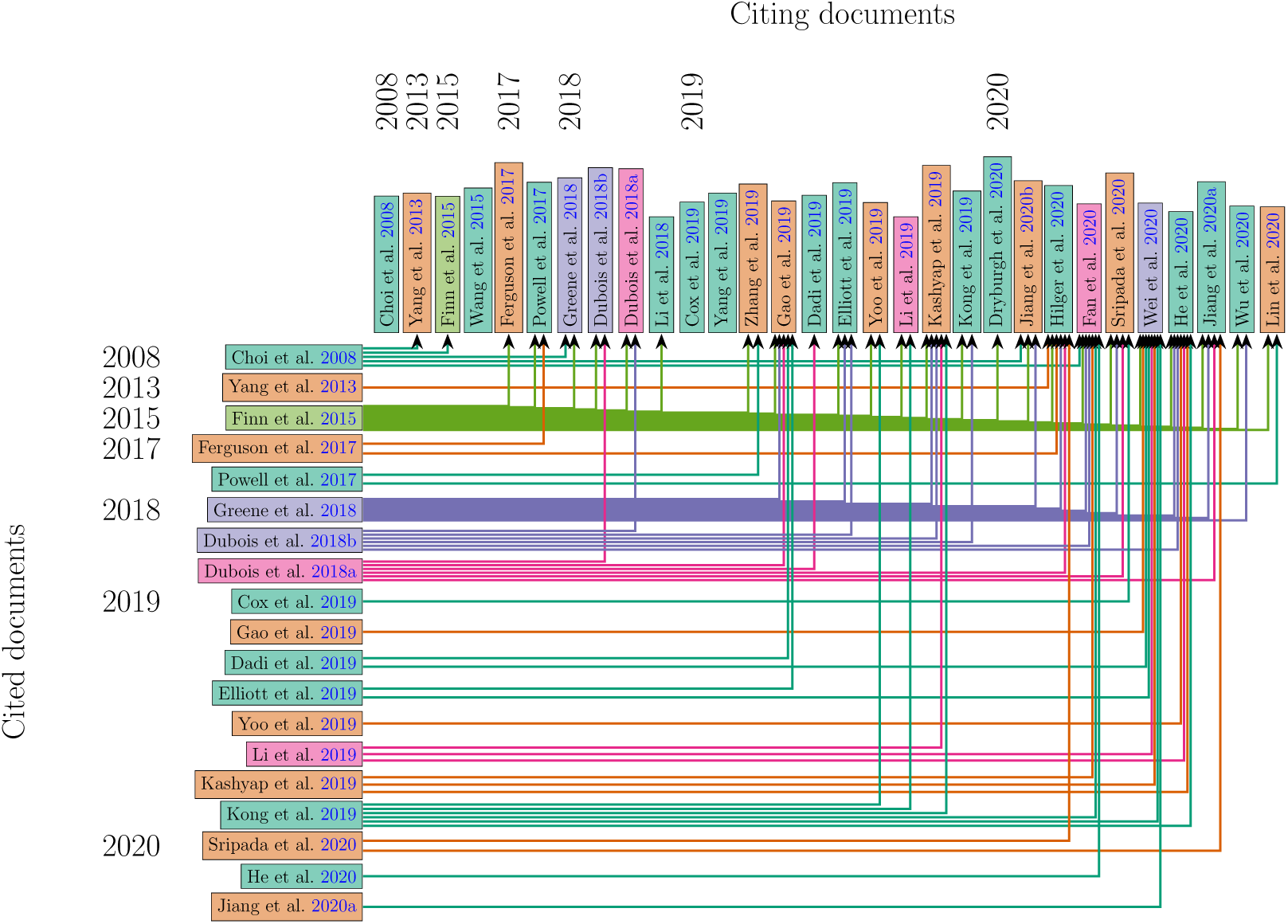
A citation network relating all 30 studies identified in Figure 1. Colors are used to better differentiate studies and carry no meaning. Arrows are colored according to parent nodes and point from the cited work to the one citing it.

Regarding data sources, 23 (77%) studies used different releases of the Human Connectome Project (HCP). Among these, 17 (57%) studies use solely HCP data. 6 (20%) studies used the HCP together with other datasets, such as the Philadelphia Neurodevelopmental Cohort (PNC) data, the Dunedin Study, Center for Biomedical Research Excellence (COBRE) and University of Electronic Science and Technology of China (UESTC), the UK Biobank, and Brain Genomics Superstruct Project (BGSP). Other sources of data included the Neuroscience Research Institute (NRI) (Choi et al. 2008; Yang et al. 2013), Korea Advanced Institute of Science and Technology (KAIST) (Choi et al. 2008), Autism Brain Imaging Data Exchange (ABIDE) (Dryburgh et al. 2020; Wang et al. 2015), UK Biobank (Cox et al. 2019), Nathan Kline Institute - Rockland Sample (NKI) (Hilger et al. 2020) and UESTC (Jiang et al. 2020b). All sources of data provide images acquired with 3 T MRI scanners, with the exception of NRI, that only includes data acquired with 1.5 T. See Figure 4a.

**Figure 4:**
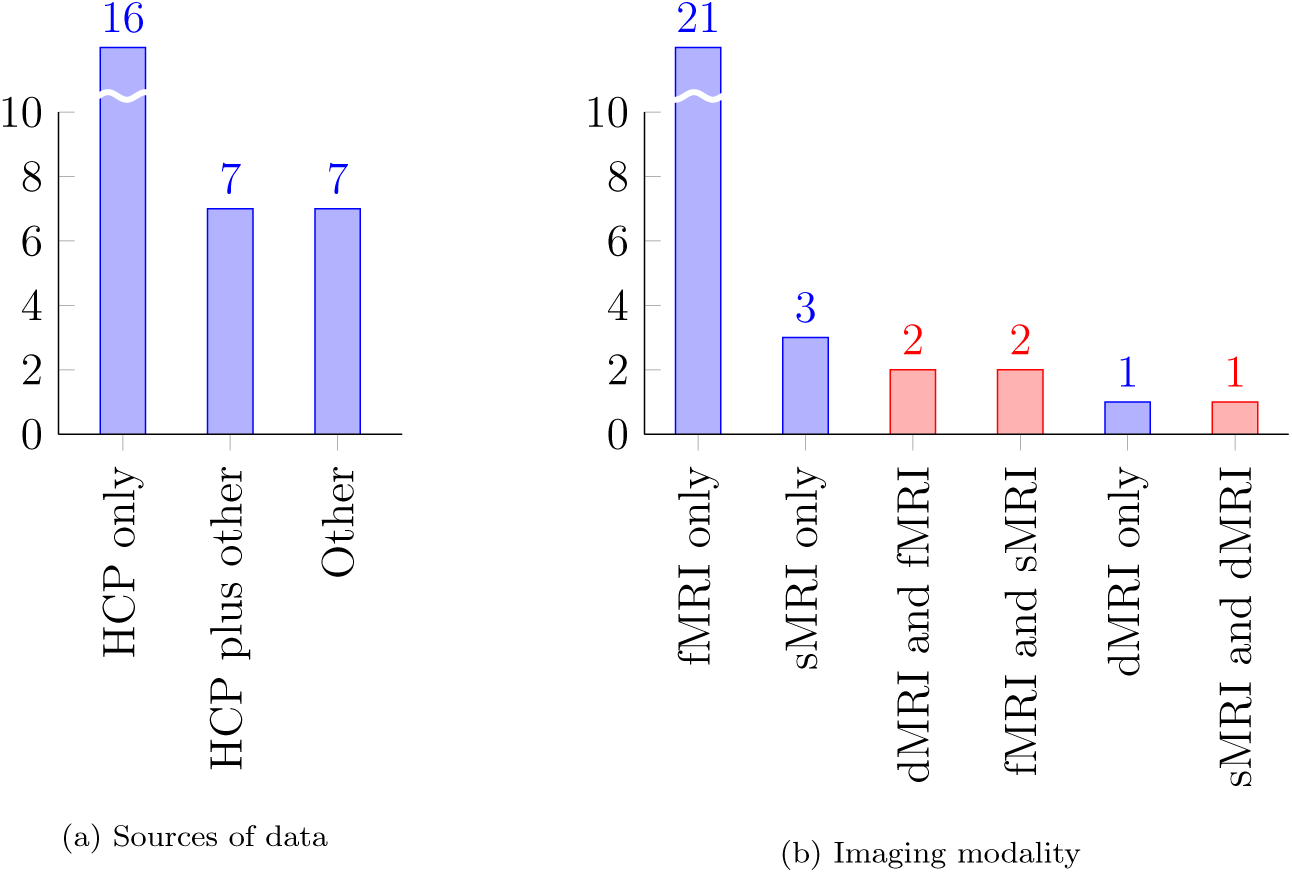
General characteristics of eligible studies. (a) shows the main sources of data identified in the sample. 23 (77%) studies employed different releases of the HCP, with 17 (57%) based solely on HCP data. (b) shows the use of different imaging modalities. Shown in blue, 25 studies were based on unimodal data: 21 (70%) used fMRI, 3 (10%) used sMRI and 1 (3%) used dMRI exclusively. Shown in red, the remaining five studies employed multimodal data: 2 (7%) used fMRI and sMRI, 1 (3%) used sMRI and dMRI, 2 (7%) used dMRI and fMRI.

Regarding imaging modality, 21 (70%) studies only used fMRI data. sMRI was the only imaging modality in 3 (10%) studies. 1 (3%) study concerned only dMRI. Multimodality was also explored, with fMRI and sMRI in 2 (7%) studies, sMRI and dMRI in 1 (3%) study, and dMRI and fMRI in 1 (3%) study. No study performed multimodal prediction based on fMRI, sMRI and dMRI simultaneously. Also, all studies used solely MRI data, *i.e.* no additional imaging such as PET, EEG or MEG is used. See Figure 4b.

We identified four constructs reported as outcomes. **G^F^** is an outcome in 20 (67%) studies, IQ in 6 (20%) studies, general intelligence, general cognitive ability or G appears in 4 (13%) studies and cognitive ability appears in 1 (3%) study. 2 (3%) studies reported results on **G^F^** and other NIH Toolbox for Assessment of Neurological and Behavioral Function (NIHTB) cognition scores (Fan et al. 2020; Wu et al. 2020), *i.e.* total, fluid and/or crystallized cognition scores. 1 (3%) study includes measures of both IQ and **G^F^** as outcomes (Jiang et al. 2020a).

The most common reported instrument is the 24-item Raven’s Progressive Matrices (RPM), appearing in 22 (73%) studies. In all these studies, the RPM employed is the Penn Matrix Test (PMAT), from the University of Pennsylvania Computerized Neurocognitive Battery (PennCNB), which also appears in 18-item format in 2 (7%) studies (Gao et al. 2019; Greene et al. 2018). The 36-item Raven’s Advanced Progressive Matrices Set II appears in 1 (3%) study (Choi et al. 2008), as one test in the estimation of G. All 20 studies that studied **G^F^** reported the usage of RPM. 3 (10%) of these studies also studied additional scores, either due to availability in specific datasets or as parallel measures. These are the Wechsler Adult Intelligence Scale (WAIS) matrix reasoning test score, as a substitute for **G^F^** in the BGSP (Li et al. 2019), and NIHTB fluid cognition scores (Fan et al. 2020; Wu et al. 2020). RPM-like tests also appear in studies that derive G from analytical decomposition of test scores (Choi et al. 2008; Dubois et al. 2018a; Sripada et al. 2020). See Table 2.

**Table 2:** On the quality of the measurement of intelligence. This categorization follows a set of rules established in Gignac et al. (2017). 1 = poor, 2 = fair, 3 = good, 4 = excellent,= unclear. FA = factor analysis; K-WAIS-R = Korean WAIS-R; WASI = Wechsler Abbreviated Scale of Intelligence; WISC-IV = Wechsler Intelligence Scale for Children - 4th edition.

| Studies | Measurement | Number of tests | Dimensions | Testing time (min) | Rating |
| --- | --- | --- | --- | --- | --- |
| Choi et al. 2008 | **G** (principal component of 36-item **RPM** and **K-WAIS-R** subtests) | 9+ | 3+ | 40+ | 4 |
| Dubois et al. 2018a; Sripada et al. 2020 | **G** (FA of 10 tests in the **NIHTB** and **PennCNB**) | 9+ | 3+ | 40+ | 4 |
| Cox et al. 2019 | **G** (FA of 4 tests in the UK Biobank) | 2–8 | 3+ | 20–39 | 3 |
| Yang et al. 2013 | **FSIQ** (**K-WAIS-R**) | 9+ | 3+ | 40+ | 4 |
| Jiang et al. 2020a | **FSIQ** (**WAIS**) | 9+ | 3+ | 40+ | 4 |
| Jiang et al. 2020a,b | **FSIQ** (Chinese **WAIS**) | 9+ | 3+ | 40+ | 4 |
| Wang et al. 2015 | **IQ** (**WISC-IV** in **ABIDE**) | 9+ | 3+ | 40+ | 4 |
| Hilger et al. 2020 | **FSIQ** (**WASI** in **NKI**) | 2–8 | 3+ | 40+ | 3 |
| Wang et al. 2015 | **IQ** (**WASI** in **ABIDE**) | 2–8 | 3+ | 40+ | 3 |
| Dryburgh et al. 2020 | **IQ** (Unclear) | ? | ? | ? | ? |
| Dadi et al. 2019; Dubois et al. 2018b; Fan et al. 2020; Ferguson et al. 2017; Finn et al. 2015; Gao et al. 2019; Greene et al. 2018; He et al. 2020; Jiang et al. 2020a; Kashyap et al. 2019; Kong et al. 2019; Li et al. 2018; Li et al. 2019; Lin et al. 2020; Powell et al. 2017; Wei et al. 2020; Wu et al. 2020; Yang et al. 2019; Yoo et al. 2019; Zhang et al. 2019 | **G<sub>F</sub>** (24-item **RPM** number of correct responses in **HCP**) | 1–2 | 1–2 | 3–19 | 2 |
| Li et al. 2019 | **G<sub>F</sub>** (**WAIS** - Matrix Reasoning test) | 1–2 | 1–2 | ? | 2 |
| Gao et al. 2019; Greene et al. 2018 | **G<sub>F</sub>** (18-item **RPM** in **PNC**) | 1–2 | 1–2 | 3–19 | 2 |
| Gao et al. 2019; Greene et al. 2018 | **G<sub>F</sub>** (24-item **RPM** in **PNC**) | 1–2 | 1–2 | 3–19 | 2 |
| Powell et al. 2017 | **G<sub>F</sub>** (24-item **RPM** total skipped items in **HCP**) | 1–2 | 1–2 | 3–19 | 2 |
| Powell et al. 2017 | **G<sub>F</sub>** (24-item **RPM** median reaction time for correct responses in **HCP**) | 1–2 | 1–2 | 3–19 | 2 |
| Wu et al. 2020 | Total cognition score (composite score from the **NIHTB**) | 2–8 | 3+ | 40+ | 3 |
| Wu et al. 2020 | Fluid cognition score (composite score from the **NIHTB**) | 2–8 | 3+ | 40+ | 3 |
| Fan et al. 2020; Wu et al. 2020 | Crystallized cognition score (composite score from the **NIHTB**) | 2–8 | 3+ | 40+ | 3 |
| Elliott et al. 2019 | Cognitive ability (**WAIS-IV** in the Dunedin study) | 9+ | 3+ | 40+ | 4 |
| Elliott et al. 2019 | Cognitive ability (24-item **RPM** in **HCP**) | 1–2 | 1–2 | 3–19 | 2 |

We used qualitative cues in titles and abstracts to determine the overall scope of studies. 10 (33%) studies had prediction of intelligence as their primary objective. Other 10 (33%) studies were concerned primarily with predictive modeling, although not focused on intelligence. 4 (13%) studies focused primarily on intelligence, but not primarily on predictive modeling. The remaining 6 (20%) studies did not focus primarily on intelligence and primarily on predictive modeling, albeit including results on both. Out of the 25 (83%) studies employing fMRI, 20 (67%) explored FC. All of these 20 include RSFC-based analyses, while 12 (40%) studied RSFC exclusively. In 19 (63%) studies, the only fMRI data was resting-state fMRI (RS-fMRI). 6 (20%) studies used T-fMRI, with task FC and/or spatial topographies as inputs (Choi et al. 2008; Elliott et al. 2019; Gao et al. 2019; Greene et al. 2018; Sripada et al. 2020; Wu et al. 2020). Choi et al. (2008) employed a fluid reasoning task. Elliott et al. (2019), Gao et al. (2019), Greene et al. (2018), Sripada et al. (2020), and Wu et al. (2020) employed seven tasks from the HCP. Additionally, Gao et al. (2019) and Greene et al. (2018) used the working-memory and emotion identification tasks from the PNC and Elliott et al. (2019) employed the emotion processing, color Stroop, monetary incentive delay and episodic memory tasks from the Dunedin study.

Not counting intracranial volume, which is used both as a predictor and as a confounder in several studies, all 6 (20%) studies reporting usage of sMRI employ morphometric measurements as predictors. The small sample of dMRI-including studies included as predictors mean diffusivity and fractional anisotropy, structural connectivity, local connectome fingerprints, and structural connectivity tensors and local structural connectivity.

Regression based on linear models was reported in 25 (83%) studies. Among these, 10 (33%) reported use of some form of penalized linear modeling. 10 (33%) reported using Connectome Predictive Modeling (CPM). 4 (13%) reported using Support Vector Regression. 5 (17%) reported using linear regression, either on inputs or on extracted components, *e.g.*, Principal Components Regression. 2 (7%) reported using Partial Least Squares Regression. Regression based on nonlinear models was reported in 5 (17%) studies. These include polynomial Kernel SVR (Wang et al. 2015), correlation kernel ridge regression (KRR) (He et al. 2020; Li et al. 2019), dice overlap KRR (Kong et al. 2019) and deep learning, based on convolutional neural networks (CNNs), graph neural networks and fully connected deep networks (He et al. 2020) or recurrent neural networks (RNNs) (Fan et al. 2020).

In 29 (97%) studies, prediction of intelligence was implemented as regression, *i.e.* prediction of a continuous variable. 1 (3%) study (Dadi et al. 2019) performed classification, subdividing subjects into two groups, one with high and the other with low IQ. They report using Support Vector Classification and Penalized Logistic Regression, as linear models, and 1-Nearest Neighbor, Näıve Bayes and Random Forest, as non-linear models.

Regarding the level of spatial abstraction of input data, 26 (87%) studies presented inputs at the regional level, either intra-regional features 7 (23%), *e.g.*, regional cortical thickness estimates, or inter-regional features in 20 (67%) studies, *e.g.*, RSFC. Inter-voxel predictors appear in 2 (7%) studies (Powell et al. 2017; Zhang et al. 2019), in the form of local dMRI structural connectivity. Intra-voxel predictors appear in 5 (17%) studies (Hilger et al. 2020; Jiang et al. 2020b; Kong et al. 2019; Li et al. 2018; Wu et al. 2020), *e.g.*, seed-based FC or voxelwise morphometry, ALFF, or T-fMRI statistical maps. No study used raw or minimally preprocessed imaging data directly as input to ML models.

In total, discounting censored and unclear results, *e.g.*, results presented only graphically, 209 results are presented across 25 studies, encompassing 10 performance metrics. These are Pearson correlation coefficient, Spearman rank correlation coefficient, R-squared, square root of R-squared, MAE, MSE, root MSE (RMSE), normalized RMSE (NRMSE), normalized root mean squared deviations (nRMSD), and area under the ROC curve (AUC). See Appendix C for the mathematical definition of each.

From the 30 studies encompassed in this review, 9 did not directly mention the tests used (Dadi et al. 2019; Dryburgh et al. 2020; He et al. 2020; Kashyap et al. 2019; Kong et al. 2019; Li et al. 2019; Wang et al. 2015; Wu et al. 2020; Yoo et al. 2019). Supplementary materials and citations were consulted to identify tests used in all but one study (Dryburgh et al. 2020). See Table 2. There was, however, little information about the measurement’s validity for the populations under study. 3 studies cited references deemed adequate (Ferguson et al. 2017; Wei et al. 2020; Yang et al. 2019), whereas partial references were cited in 2 studies (Hilger et al. 2020; Lin et al. 2020). Regarding measurement quality, 7 measurements were rated as excellent, distributed across 8 (27%) studies. 6 measurements were rated as good, distributed across 5 (17%) studies. 7 measurements were rated as fair, distributed across 21 (70%) studies. 1 (3%) study has a measurement of IQ which we could not identify, based on pre-processed ABIDE, which include multiple instruments. See Table 2 for detailed ratings.

The assessments of RoB and applicability are shown in Table 3. In total, 8 (27%) studies were rated with low overall RoB and low concern regarding applicability. This includes five development-only studies (Dubois et al. 2018a,b; He et al. 2020; Li et al. 2019; Sripada et al. 2020), one development-validation study (Cox et al. 2019), and the validation portions of two development-validation studies (Elliott et al. 2019; Greene et al. 2018). These are eligible for quantitative synthesis, *i.e.* meta-analysis. Li et al. (2019) does not present prediction results in text format however, and thus was not used. Results pertaining to sMRI and dMRI encompass only the 4 results in Cox et al. (2019), and thus these modalities were ineligible for quantitative synthesis, per our protocol. 89 results identified among the remaining 6 studies were suitable for quantitative synthesis: 3 in Dubois et al. (2018a), 8 in He et al. (2020), 16 in Sripada et al. (2020), 39 in Dubois et al. (2018b), 6 in Greene et al. (2018), and 17 in Elliott et al. (2019). All of these employed fMRI solely and reported either the Pearson Correlation Coefficient or R-squared, with the exception of Greene et al. (2018), which reported squared Spearman Rank Correlation. We opted to group this result with R-squared.

**Table 3:**
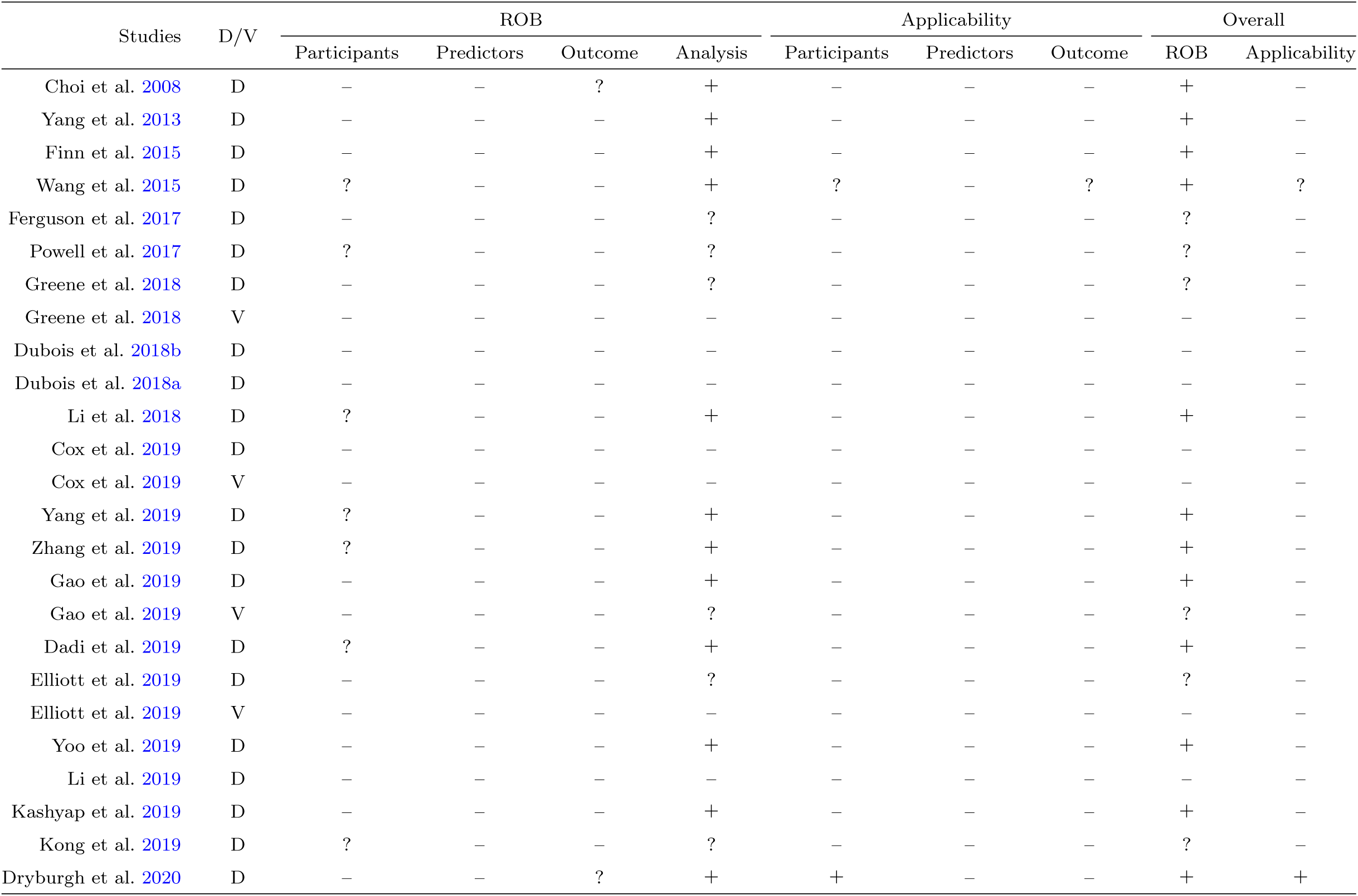
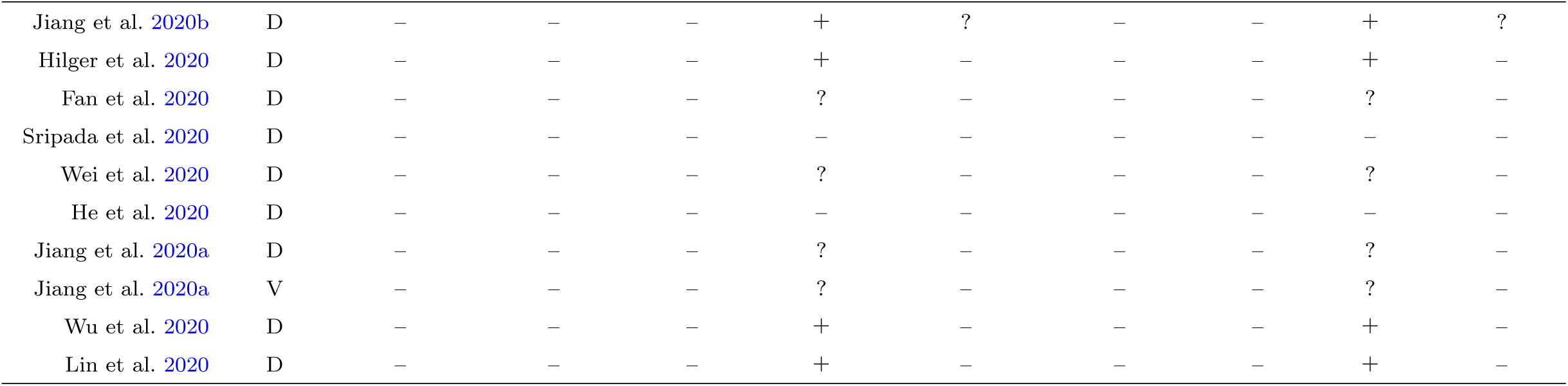
PROBAST = Prediction model Risk Of Bias ASsessment Tool; ROB = risk of bias; D = Development; V = Validation. + expresses low ROB/low concern regarding applicability; – expresses high ROB/high concern regarding applicability; andexpresses unclear ROB/unclear concern regarding applicability.

| Studies | D/V | ROB |  |  |  | Applicability |  |  | Overall |  |
| --- | --- | --- | --- | --- | --- | --- | --- | --- | --- | --- |
|  |  | Participants | Predictors | Outcome | Analysis | Participants | Predictors | Outcome | ROB | Applicability |
| Choi et al. 2008 | D | – | – | ? | + | – | – | – | + | – |
| Yang et al. 2013 | D | – | – | – | + | – | – | – | + | – |
| Finn et al. 2015 | D | – | – | – | + | – | – | – | + | – |
| Wang et al. 2015 | D | ? | – | – | + | ? | – | ? | + | ? |
| Ferguson et al. 2017 | D | – | – | – | ? | – | – | – | ? | – |
| Powell et al. 2017 | D | ? | – | – | ? | – | – | – | ? | – |
| Greene et al. 2018 | D | – | – | – | ? | – | – | – | ? | – |
| Greene et al. 2018 | V | – | – | – | – | – | – | – | – | – |
| Dubois et al. 2018b | D | – | – | – | – | – | – | – | – | – |
| Dubois et al. 2018a | D | – | – | – | – | – | – | – | – | – |
| Li et al. 2018 | D | ? | – | – | + | – | – | – | + | – |
| Cox et al. 2019 | D | – | – | – | – | – | – | – | – | – |
| Cox et al. 2019 | V | – | – | – | – | – | – | – | – | – |
| Yang et al. 2019 | D | ? | – | – | + | – | – | – | + | – |
| Zhang et al. 2019 | D | ? | – | – | + | – | – | – | + | – |
| Gao et al. 2019 | D | – | – | – | + | – | – | – | + | – |
| Gao et al. 2019 | V | – | – | – | ? | – | – | – | ? | – |
| Dadi et al. 2019 | D | ? | – | – | + | – | – | – | + | – |
| Elliott et al. 2019 | D | – | – | – | ? | – | – | – | ? | – |
| Elliott et al. 2019 | V | – | – | – | – | – | – | – | – | – |
| Yoo et al. 2019 | D | – | – | – | + | – | – | – | + | – |
| Li et al. 2019 | D | – | – | – | – | – | – | – | – | – |
| Kashyap et al. 2019 | D | – | – | – | + | – | – | – | + | – |
| Kong et al. 2019 | D | ? | – | – | ? | – | – | – | ? | – |
| Dryburgh et al. 2020 | D | – | – | ? | + | + | – | – | + | + |

Table 3: PROBAST = Prediction model Risk Of Bias ASsessment Tool; ROB = risk of bias; D = Development; V = Validation.
| Studies | D/V | ROB |  |  |  | Applicability |  |  | Overall |  |
| --- | --- | --- | --- | --- | --- | --- | --- | --- | --- | --- |
|  |  | Participants | Predictors | Outcome | Analysis | Participants | Predictors | Outcome | ROB | Applicability |
| Jiang et al. 2020b | D | – | – | – | + | ? | – | – | + | ? |
| Hilger et al. 2020 | D | – | – | – | + | – | – | – | + | – |
| Fan et al. 2020 | D | – | – | – | ? | – | – | – | ? | – |
| Sripada et al. 2020 | D | – | – | – | – | – | – | – | – | – |
| Wei et al. 2020 | D | – | – | – | ? | – | – | – | ? | – |
| He et al. 2020 | D | – | – | – | – | – | – | – | – | – |
| Jiang et al. 2020a | D | – | – | – | ? | – | – | – | ? | – |
| Jiang et al. 2020a | V | – | – | – | ? | – | – | – | ? | – |
| Wu et al. 2020 | D | – | – | – | + | – | – | – | + | – |
| Lin et al. 2020 | D | – | – | – | + | – | – | – | + | – |

Forest plots with individual results are shown in Appendix E. For the Correlation coefficient obtained from fMRI, both G and **G^F^** have expected correlations significantly different from zero, based on 66 results from 5 studies (Dubois et al. 2018a,b; Elliott et al. 2019; He et al. 2020; Sripada et al. 2020). For G, the expected correlation was 0.42 (CI_95%_ = [0.35, 0.50], p *<*0.001). For **G^F^**, the expected correlation was 0.15 (CI_95%_ = [0.13, 0.17], p *<*0.001). Both are significantly different (p *<*0.001). A significant difference between HCP and UK Biobank was found: 0.086 (CI_95%_ = [0.012, 0.16], p = 0.022). Residual heterogeneity was estimated at *I*^2^= 77.8% for this analysis. For R-squared, only G has expected R-squared significantly different from zero, based on 34 results from 6 studies (Dubois et al. 2018a,b; Elliott et al. 2019; Greene et al. 2018; He et al. 2020; Sripada et al. 2020). For G, the expected R-squared was 0.16 (CI_95%_ = [0.13, 0.18], p *<*0.001). For **G^F^**, the expected R-squared was 0.022 (CI_95%_ = [-0.021, 0.066], p = 0.3). Both are significantly different (p *<*0.001). No significant differences between HCP and PNC or between HCP and UK Biobank were found. Residual heterogeneity was estimated at *I*^2^= 63.3% for this analysis.

TRIPOD has items that apply only to either validation or development of models. Here, all studies included development of models, while a few also included external validation. We chose to represent results together in Figure 5, with the caveat that a few items (10e, 12, 13c, 17, 19a) only apply to studies that include validation of models.

**Figure 5:**
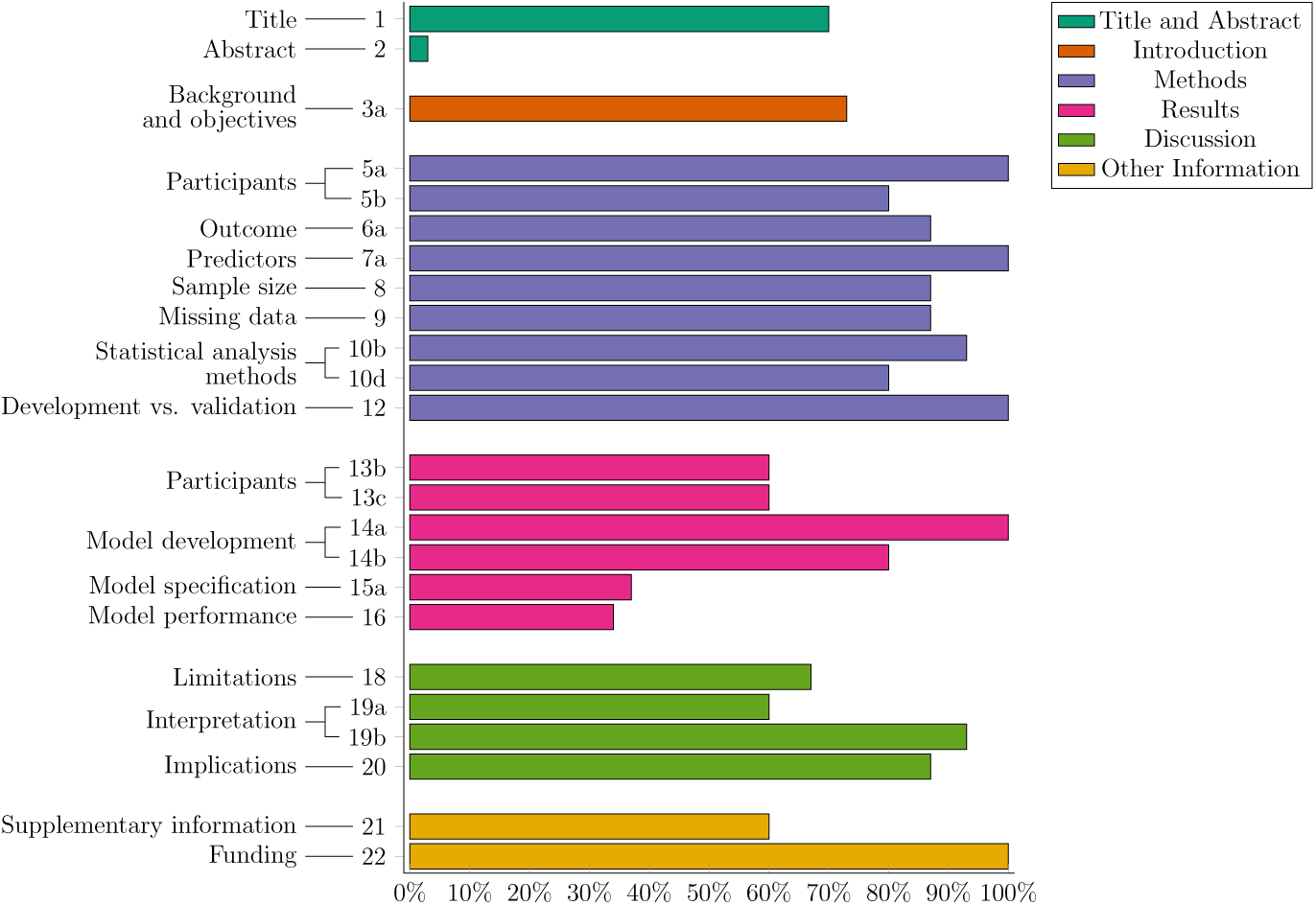
Overall results from the TRIPOD assessment of reporting quality. Bars represent average scores across studies. Items are nested into topics which are nested within sections, following the specification in TRIPOD. Sections and topics are shown, while items can be inspected in more detail in Table D.5 or Heus et al. (2019) and Moons et al. (2015). Items 7a, 10b and 15a were adjusted following Wang et al. (2015). Table D.5 reflects these adjustments.

The histogram of TRIPOD ratings is shown in Figure 6.

**Figure 6:**
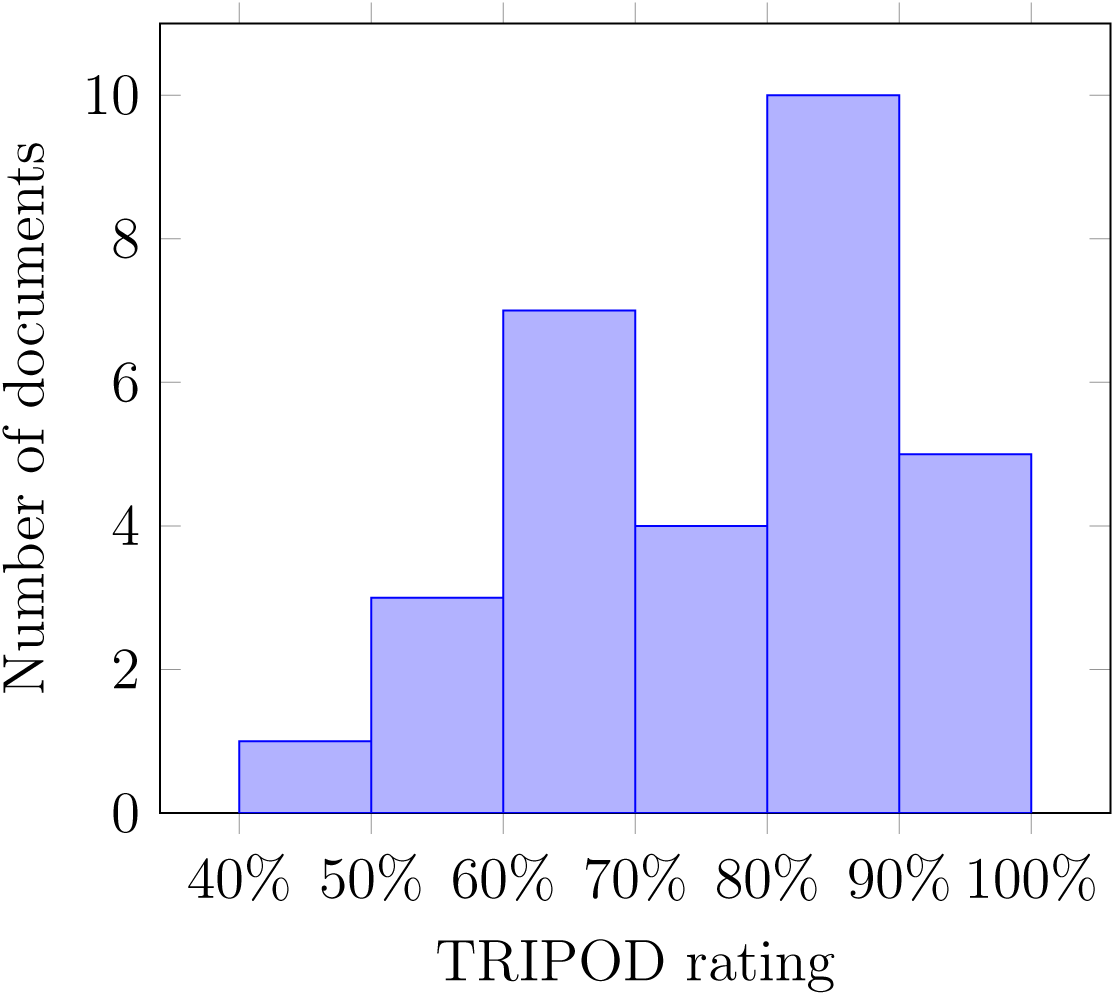
Distribution of TRIPOD overall ratings across 30 studies.

Funnel plots for both the analysis of correlation coefficients and R-squared are shown in Appendix F. Both analyses present symmetrical funnel plots, which imply low risk of publication bias, but the range of standard errors is low, due to sample limitations, *e.g.* the lack of results with more subjects.

We additionally analyzed the relationship between the expected effect size and training set size. Due to the small number of results pertaining to R-squared, this analysis was performed only for the correlation coefficient. Figure 7 shows the expected correlation coefficient between predicted values and true labels as a function of approximate training set size. This comparison is qualitative, and does not take into account confounders, but it is also expected that such procedures are more robust in larger sample sizes. Compare with Figure E.8, which includes only studies with low RoB and low concerns regarding applicability.

**Figure 7:**
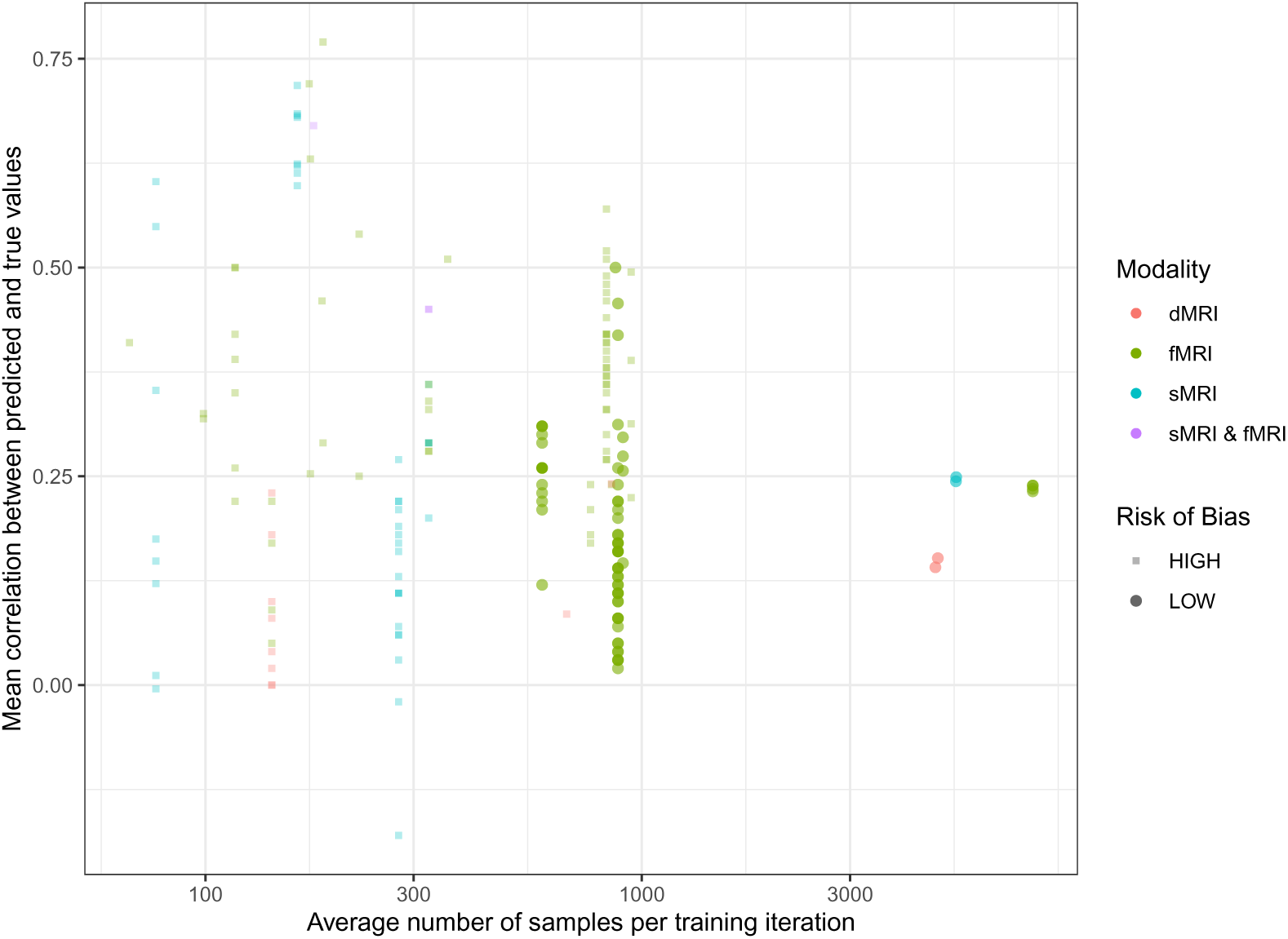
The expected correlation coefficient according to the approximate training size data employed across studies. With the exception of holdout-based studies, where the actual training set size is known, the approximate training size data was estimated as the total number of data available for training times by (*K −* 1)*/K*, where *K* is the number of groupings. Low risk of bias refers to studies that were rated with low RoB and low concern regarding applicability in Table 3. Modality refers to the imaging modality of each individual result.

## 4. Discussion & Conclusion

Here, we systematically reviewed available studies on the application of ML to the prediction of human intelligence using MRI data. Most of these studies were published very recently. See Figure 2. Namely, two-thirds were published in 2019 and 2020. This attests the high and growing interest over this question in the literature.

It is also very clear from Figure 3 that some highly cited studies exert a larger influence in the literature. Later works were highly influenced by these and, in a way, the current state of the literature reflects those earlier successes. A few studies do not cite other earlier studies in Figure 3, likely because not every document focused exclusively on individualized prediction and/or intelligence. That should be taken into account when examining most results, especially TRIPOD ratings. See the TRIPOD checklist (Heus et al. 2019).

In the case of T-fMRI, results are largely compatible across datasets, but not across tasks. Gao et al. (2019) and Greene et al. (2018) show that FC derived from tasks are stronger predictors of **G^F^** than RSFC. The gambling and the working-memory tasks demonstrate higher predictive power. Sripada et al. (2020) and Wu et al. (2020) also found that the working-memory task is highly discriminative of G, this time using statistical spatial maps.

While some of the studies presented results on more than one MRI modality, only one study presented a model that learns from multimodal data. Jiang et al. (2020b) presented results on both vertexwise cortical thickness and region of interest (ROI)-based RSFC. They show that a model that uses both modalities at once attains significantly higher predictive accuracy for intelligence compared to single-modality models. Choi et al. (2008) “neurometric model” includes both cortical thickness and T-fMRI statistical maps as inputs, but each part of the model was learned in isolation.

The HCP (Essen et al. 2013; Glasser et al. 2016) is the most employed dataset, appearing in 73% of the sample. Dating its first releases back to 2013, it began being employed for the prediction of intelligence as early as 2015 (Finn et al. 2015).

The majority, encompassing 83% of eligible studies, employed linear modeling for regression to some extent. Linear modeling is a strong baseline, also appearing in studies employing non-linear models. The most popular linear approaches include CPM and penalized linear models, each appearing in 40% of studies using linear models. CPM (Shen et al. 2017) is a very streamlined approach to predictive modeling. It is based on building linear models to predict outputs from aggregate measures of correlation between inputs and outputs, after thresholding based on significance. Features that are kept are then divided into positive-feature and negative-feature networks (Finn et al. 2015). Features in each network are summarized, *e.g.*, summed or averaged, for each sample. Then, linear regression is used to predict outputs from these aggregate features, either separately or jointly for the positive-feature and negative-feature networks. After its introduction by Finn et al. (2015), albeit not yet named CPM, it became a staple of predictive modeling. Even though “connectome” appears in its name, the same principle can also be extended to other domains such as morphometry (Jiang et al. 2020b). Penalized linear modeling, on the other hand, does not aggregate features. Often, univariate filtering based on significance thresholding is used, akin to CPM. Then, however, remaining features are used as they are, without any additional transformation. The rationale for it is that penalization of coefficients can resolve commonalities and differences in features, and effectively attenuates overfitting.

Non-linear regression modeling appears in only few studies, 17% of the sample. This might be due to the intrinsic high dimensionality of neuroimaging data, particularly evident for fMRI. At such high dimensionality, overfitting becomes a greater concern for more flexible models. The only non-linear model appearing more than once is KRR, a kernelized penalized linear regression. It is a very flexible approach given that a similarity measure between samples can be derived. Instead of using the base features in the model, features are expanded to higher (potentially infinite) dimensionalities. The kernel is the dot product between samples in this high dimensional space, which allows for efficient computation of models, bypassing the need of explicitly computing features in the new basis. In the sample, the correlation and the Dice overlap kernels were used in different studies. Due to the implicit high dimensionality, penalization is used very often, such as the ridge penalty, in the case of KRR.

Across studies, prediction is usually performed in aggregate measures of the data. Abrol et al. (2021) systematically shows that deep neural networks when trained on raw data outperform classical linear and non-linear ML models in the prediction of age, gender and Mini Mental State Examination scores. They also show that embeddings obtained from deep neural networks provide strong features for classical ML. This suggests that the choice of features in the literature has the potential of negatively biasing the performance of deep neural networks. Deep neural networks allow for using structured data, due to their inductive biases, present in architectures such as CNNs for image data or RNNs for sequence data. Only few studies use deep neural networks for the prediction of intelligence using neuroimaging. He et al. (2020) modeled **G^F^** based on RSFC with three deep neural networks. Fan et al. (2020) modeled **G^F^** and **G^C^** based on dynFC with RNNs. Vieira et al. (2021b, not in this review) implements prediction of G also with RNNs, but based on RS-fMRI timeseries.

### 4.1. Limitations across studies

We must first state that limitations found in the analyzed studies have to be examined under the light of the current review’s question, *i.e.* what current literature regarding the ML-based prediction of intelligence using neuroimaging looks like. Many studies did not focus primarily on the prediction of intelligence, even though they included such results. Studies proposing or benchmarking modeling choices, *i.e.* preprocessing, ML, and imaging methods, will often include intelligence among their results. A common occurrence in these studies, that include several outcomes, is that they will not give results in text format. When results are shown only graphically, we decided to not use inferred numbers. Also, when assessing the TRIPOD checklist, we only scored items that were clearly within the scope of the document. For example, studies not primarily concerned with prediction were not penalized by not mentioning prediction in their title, *i.e.* item 1.ii in TRIPOD.

The choice of outcome may also be an object of discussion. Lohman et al. (2012) argue that **G^F^** consists of three components: sequential reasoning, quantitative reasoning and inductive reasoning. The latter is the core of RPM. For this reason, Gignac (2015) argues that RPM can be considered an imperfect measure of **G^F^**. This is due to its rather narrow scope, since it exclusively consists of figural type items. All studies that predicted **G^F^** employed the RPM in some extent. Most, 19 out of 20, used solely the RPM, with the remaining one employing both the RPM and NIHTB’s fluid composite score. For this reason, their results necessitate further consideration.

For both correlation and R-squared results, G-based results are significantly higher than **G^F^**-based ones. This alludes to Gignac et al. (2017), who showed that higher measurement quality moderates the observed correlation between intelligence and brain volume. In our assessment in Table 2, G derived from 10-tests in the NIHTB and PennCNB was rated as excellent, while **G^F^** or “cognitive ability” obtained from a single test was rated as fair. Furthermore, Dubois et al. (2018a,b) used the same predictor data based on RS-fMRI, but obtained very disparate results using the HCP. The authors reported *r* = 0.263 and *R*^2^ = 0.047, when predicting PMAT-based **G^F^**, versus *r* = 0.457 and *R*^2^ = 0.206, when predicting G based on the factor analysis of 10 tests in the PennCNB and NIHTB.

The measurement of G and **G^F^** may incur risks of bias compromising proper estimation of intelligence. The use of a single-domain test, such as inductive reasoning in RPM, would evaluate an isolated skill and not measure adequately intelligence, which is by definition a set of different cognitive skills. Furthermore, a test that assesses different skills needs to cover more than one cognitive domain, *e.g.*, verbal, visual or spatial, to obtain a complete measurement (Gignac et al. 2017).

Another bias in interpreting results can occur due to the omission of information related to the measurement of intelligence. The article must present the construct, *e.g.* intelligence, G, or **G^F^**, and the psychological test used so that it is possible to verify whether the test is adequate to measure the function contained in the specific construct. However, a psychological test suitable for the construct is not necessarily suitable for the population studied. It is essential to ensure tests are adequately validated for the population under study.

Despite the solid empirical basis of the concepts of G, **G^F^**, and **G^C^**, there are still concerns regarding cognitive abilities associated with G (Kent 2017). New research on the neuroimaging-based prediction of intelligence should bring more specifications when evaluating cognitive constructs, such as the psychological instrument, validity, and application range.

The current conceptualization of the intelligence construct does not encompass only **G^C^** or **G^F^**. It covers adaptability and problem-solving in real life, considering emotional intelligence factors, decision making (Stankov 2017), and personality (Kent 2017). The interaction of these cognitive processes in an integrated way configures a complex multidimensional construct (McGrew 2009). Due to this characteristic, it is recommended to use as many specifications as possible when performing the intelligence measurement.

The best model for the development of psychological instruments in intelligence evaluation is the Cattell-Horn-Carroll (CHC), seen as the best psychometric evidence for human aptitudes (Abu-Hamour et al. 2016; Hurks et al. 2016; James et al. 2015; Lecerf et al. 2010; Wechsler et al. 2016). CHC theory consists of a hierarchical multidimensional model with ten factors of cognitive functioning: Fluid intelligence (Gf), Quantitative knowledge (Gq), Crystallized intelligence (Gc), Reading and writing (Grw), Short-term memory (Gsm), Visual processing (Gv), Auditory Processing (Ga), Long-term memory storage and retrieval ability (Glr), Processing Speed (Gs) and Decision speed (Gt). However, there is criticism over its weak explanatory capacity, its failure to make testable predictions, and its enmeshment to the Woodcock-Johnson battery (Wasserman 2019). The Woodcock-Johnson battery of tests (Woodcock et al. 2001) was designed to be more aligned to the CHC theory. However, there is evidence against this alignment and the lack of support for interpreting most of the scores suggested by its scoring system (Dombrowski et al. 2019). To date, no psychological test measures the broad cognitive abilities established in the CHC model which are contained in intelligence. For an adequate measurement, one should make use of instruments that are most related to the CHC theory, *e.g.*, WAIS or Woodcock-Johnson Tests, Fourth Edition (WJ IV ACH).

The preponderance of **G^F^** has three probable causes: (1) early success, as reported in Finn et al. (2015), which is cited by 23 out of 26 possible studies, as can be seen in Figure 3; (2) ease of estimation, since it is often taken to comprise the score of a single test; and (3) availability, which compounds with the last reason, since RPM scores are available from the HCP, UK Biobank, BGSP and PNC.

The prevalence of **G^F^** presents some challenges regarding the validity of results. The RPM can be considered a good score to include for the estimation of G and **G^F^**. Current studies show that **G^F^** and G have a strong correlation and are often statistically indistinguishable (Caemmerer et al. 2020). In isolation, however, according to the criteria published in Gignac et al. (2017), the RPM would be considered at best a “fair” measure of G. Similarly, although it is correlated with **G^F^**, it does not appear to be remarkable in comparison with other tests that measure **G^F^** (Gignac 2015). These findings point to the necessity of investigating what the models are learning through the RPM, and how much of it is shared between G, **G^F^** and test specific variance. This would better clarify how much the prediction of RPM correlates with prediction of **G^F^**.

The literature constructs a clear picture regarding the level of expected evidence: correlations between brain imaging data and intelligence are substantial, albeit reliably low. It hovers around between 0.12 and 0.25 in large sample-size studies based on the UK Biobank (Cox et al. 2019; He et al. 2020), shown in Figure 7. According to our quantitative analysis, the confidence interval covers between 0.35 and 0.50 for G and 0.13 and 0.17 for **G^F^**based on fMRI data only. A possible explanation for this is that, in fact, the current data only affords such a level of performance. This also means that unexplained components of intelligence could be potentially learned in other spatio-temporal resolutions and imaging contrast mechanisms. Another, more problematic hypothesis is that ML is capturing relationships with other behaviors and demographics that correlate with intelligence, but not intelligence itself. This “shortcut learning” (Geirhos et al. 2020) is a major challenge for ML generalizability and interpretability. Possible shortcuts could include attention and arousal, but can go much deeper, to include substance abuse, malnutrition or socioeconomic status. Population modelling is one alternative to estimate how much brain data contributes to prediction of mental traits, *i.e.* Dadi et al. (2021, not in this review) demonstrates that, despite statistical significance, multimodal brain data constributes little to the prediction of **G^F^** compared with sociodemographics.

On the choice of performance metrics, we see that Pearson correlation coefficient and R-squared are the most common in the literature. This is due to their scale invariance and perceived ease of interpretation. Despite their popularity, both are prone to biases. The correlation coefficient represents the linear association between predictions and true outcomes. Its formulation does not involve actual residuals, so models with arbitrarily large errors can still achieve perfect unitary correlation. Since R-squared involves a ratio, the denominator that represents the variance of true values can arbitrarily reduce or augment it. In other words, too small (or too large) variance of intelligence in the sample can lead to small (or large) R-squared, even under the same model (Alexander et al. 2015). This means that comparisons between studies, specially when their outcomes and/or populations differ, is at elevated risk of bias. A different choice of population incurs different characteristics of the outcome variance, possibly compromising the comparison. Model comparison on the same data could be performed under a well-behaved metric, such as the MSE or MAE.

We detected censoring for studies with high RoB and small sample sizes, as can be seen in Figure 7. Their variability and the frequency of negative results diminish with models trained on less than 300 subjects. This is a qualitative indicator of publication bias, but also of selective reporting, since most studies report comparisons with multiple models. This selective reporting can be a result of the issue described in Hosseini et al. (2020), where authors perform optimization of their models on the same data that performance is measured, leading to inflated performance estimates due to overfitting to the test set and leakage.

The diversity of populations under study across studies is skewed towards a select group of countries. The 11 datasets identified can be grouped accordingly into United States (HCP, NKI, PNC, BGSP, COBRE), New Zealand (the Dunedin Study), United Kingdom (UK Biobank), China (UESTC), South Korea (NRI, KAIST) and North America/Europe (ABIDE-I). Earlier releases of ABIDE were for the most part based on United States populations as well (New York University, Kennedy Krieger Institute, Stanford, Oregon Health & Science University, University of California, Los Angeles as in Wang et al. (2015)). This limitation stems from economic factors that affect countries differently. While some datasets sampled highly-educated young adult populations, several others are matched samples from the local general population, which diminishes risks of biases. The prediction of **G^F^** from the HCP, specially that assessed by the RPM, is very predominant in the literature. Albeit large datasets are often employed, the homogeneities across studies raise concerns regarding generalizability to other populations. Future works could perform validation analyses of trained models on new datasets, taking special care of differences in imaging acquisition and pre-processing.

While earlier association works helped to foment new theories on intelligence, current ML-based works have not yet contributed substantially to this endeavor. This comes from the fact that the majority of the works do not try to extract explanatory value from the trained models. Few works test the leverage of different features and how these fit within or without theories such as Parieto Frontal Integration Theory (P-FIT) and Network Neuroscience Theory. Future works and possibly meta-analyses can solidify these findings, providing support for existing or new theories.

A common occurrence in the assessment of PROBAST was that studies did not take into account the optimistic bias of confounders received high RoB ratings for “Analysis” Table 3. A notorious confounder which should be taken into account is kinship, in datasets like the HCP (Dubois et al. 2018a). Other, more pervasive ones, include movement and brain volume, but also sex and age. Two common approaches in the literature are removing the effect of confounders using linear models and stratifying data in a way to minimize bias due to confounders, the latter a very common approach when dealing with family structure. While our work cannot determine optimal strategies for treatment of confounders, low RoB studies were expected to recognize their effects and account for it in results.

Another common factor leading to high RoB was small sample size. It is a well-known fact from the literature that ML-based studies suffer spurious correlation induced by small samples (Varoquaux 2018). The few studies that report the standard error of the mean cross-validated performance also likely underestimate it (Varoquaux 2018). Recognizing the negative impact of small sample sizes, having fewer than 500 subjects was considered as an indicator of possible RoB in the assessment of Table 3.

### 4.2. Limitations of the review

Some possible limitations can be identified in our review methodology. Searching for manuscripts on predictive modeling on neuroimaging is particularly challenging. In the early literature, the term “predict” would often be used to refer to studies on correlations and associations. For this reason, we had to use a search strategy based on domain-knowledge. This choice, however, incurs the risk of selection bias due to missing documents. Since we successfully retrieved a reasonable number of documents, we believe that we minimized this risk and also obtained a representative sample. It is however expected that our selection missed documents, but we believe that this number should be small.

The fact that most studies either did not focus solely on intelligence or were not primarily about individualized prediction makes data retrieval difficult. For this reason, in several instances constructs and instruments are not readily identified in searchable text. We thoroughly searched for information in actual figures and supplementary materials. We did not follow citations or other sources to infer this information, since the construct should ideally be stated by authors.

Another source of variance is the fact that terminology is flexible. Studies will often use terms like cognitive ability or others with ambiguous meaning. For example, in Elliott et al. (2019) “cognitive ability” refers to both FSIQ and **G^F^**, while in Sripada et al. (2020) “general cognitive ability” names a measurement that is identified with G in other studies. Some works will refer to a G-like construct as general intelligence, others will refrain from using the term intelligence altogether. We tried to disambiguate authors’ choices with the coherence of the review in mind. This is particularly evident in Table 2, where we tried to unify terminology.

We adopted the TRIPOD adherence assessment form (Heus et al. 2019) to evaluate reporting quality. That benefits objectivity in this analysis. Measuring adherence to a specific reporting guideline has the disadvantage of potentially misrepresenting studies. This guideline is not enforced by journals, reviewers or the authors themselves in this research area. This form has been similarly applied to documents published prior to TRIPOD (Zamanipoor Najafabadi et al. 2020). Due to the generality of TRIPOD items, we believe that the risk of bias is low regarding the assessment of reporting quality. Several studies achieved high ratings, as can be seen in Figure 6.

We employed PROBAST to assess RoB and applicability. PROBAST is a tool designed primarily for studies in health and medicine, but its items are still very applicable to our review question. Another benefit is that the use of standardized tools minimizes biases when compared with an alternative created by authors. This was a post-hoc adaptation from the protocol in Vieira et al. (2021a), but, with aforementioned justification, we also consider that the risk of inducing bias is low.

We employed the PRISMA checklist as a reporting guideline. PRISMA was designed for studies that evaluate healthcare interventions, but most items can be applied to our review question. We believe that this choice offers no additional risk of bias for our review.

The quality of measurement of intelligence was evaluated by the first three criteria of the essential guide for categorizing the quality of general intelligence measurement (Gignac et al. 2017). Although the guide was proposed for G, we also used it to assess the quality of measurement of **G^F^**.

The number of studies using modalities other than fMRI with low RoB and low concerns regarding applicability was insufficient for quantitative analysis. For this reason, we only obtained meta-estimates of correlation and R-squared from fMRI. Figure 7 seems to point towards an approximately unique ceiling in performance, but the small number of studies, especially truly multimodal ones, makes that inference inconclusive.

### 4.3. Future work

Future work could explore other imaging techniques, such as PET, EEG and MEG. These imaging techniques probe different functional aspects from fMRI. PET allows the study of slow metabolic dynamics in the brain and was fundamental for the definition of the P-FIT, being employed in the study of metabolic response differences under cognitively demanding tasks (Jung et al. 2007). EEG and MEG, on the other hand, probe fast electrical cerebral dynamics, and their importance was also acknowledged in P-FIT, albeit neither was part of its experimental foundation. In addition to other imaging techniques, multimodality presents an avenue for future research. It is currently not possible to establish whether information learned from different modalities overlap due to the lack of large numbers of multimodal models. Studies employing two or more techniques or modalities at once can better disambiguate the predictive power exclusive to each. This type of study is, however, becoming more widespread. Jiang et al. (2020b) model anatomical and RSFC data jointly, Dhamala et al. (2021, not in this review) use dMRI structural connectivity and RSFC, and Dadi et al. (2021, not in this review) includes joint modeling based on RSFC, dMRI diffusion measurements, and sMRI global and regional volumes.

Most works employ ROI-level features. Although this “summarization” makes ML more amenable, since it diminishes the dimensionality of data, this level of spatial abstraction can discard useful intra-regional information. Feilong et al. (2021, not in this review) systematically demonstrates that accounting for fine-grained, intra-ROI task and resting-state FC differences lead to improvements in the prediction of G and other intelligence measurements. Future developments on data-efficient ML models that can robustly learn from minimally preprocessed data have the potential of resolving this abstraction and discovering relationships hidden by summarization.

Other ML algorithmic developments can improve prediction accuracy and validity in the future. In particular, interpretable and explainable models can further corroborate, falsify and augment current theories on the biological bases of intelligence, which were majoritarily developed based on coarse-grained spatial attributes of brain anatomy and function.

Refinements of psychometric and neuroscientific theories of intelligence will also lead to a demand for future work. Intelligence differences do not occur in isolation, being permeated by other human behaviors and environmental factors. The extended P-FIT (ExtPFIT) was formulated in Gur et al. (2021), and its generalizability can be tested in a ML-based framework. Other neuroscientific theories and extensions will probably emerge in the future.

Finally, larger scale datasets will diminish small sample-size biases in predictive models (Varoquaux 2018). Jointly learning across different datasets and discarding confounding information efficiently can boost predictive accuracy. Future works can help answer if the patterns observed in current models generalize across different populations, socio-economic environments, languages and cultures.

### 4.4. Conclusions

More than half of the identified studies include linear modeling to predict RPM-based **G^F^** from HCP fMRIs. This fact attests the significance and reliability of fMRI-based prediction studies. It also alludes to possible new avenues of research that have been studied infrequently if at all.

By pointing out salient results across studies and limitations, we hope that this work contributes to further developments in this area of research. While predictive modeling “best-practices” are abound, the literature currently lacks reporting guidelines, which could be fulfilled to ease literature search. Some gaps that can be filled by future studies include: extending and validating the current models in new populations, developing models using other spatiotemporal resolutions, other modalities, and imaging techniques, and disambiguating the contribution of neuronal phenomena to the predictions.

## Acknowledgements

Authors acknowledge funding received through grant #2018/11881-1, São Paulo Research Foundation (FAPESP) (B.H.V - PhD scholarship).

## Disclosure Statement

Authors declare the following that could be perceived as a conflict of interest: authors B.H.V. and C.E.G.S. maintain active collaboration with J.D. (Dubois et al. 2018a,b) and V.D.C. (Jiang et al. 2020a,b). J.D. and V.D.C. were not consulted or informed in any form regarding the current study.

## Appendix A. SCOPUS search string

( TITLE-ABS-KEY ( ( “cortical thickness“ OR “functional connectivity“ OR “structural connectivity“ OR “effective connectivity“ OR mri OR fmri OR morphometry ) AND ( prediction OR predict OR cpm OR “multivariate pattern analysis“ OR bases OR variability OR mvpa ) ) AND ( TITLE ( intelligence OR behavioral OR behavior OR “cognitive ability“ ) ) )

## Appendix B. PRISMA 2009 Checklist

**Table B.4:**
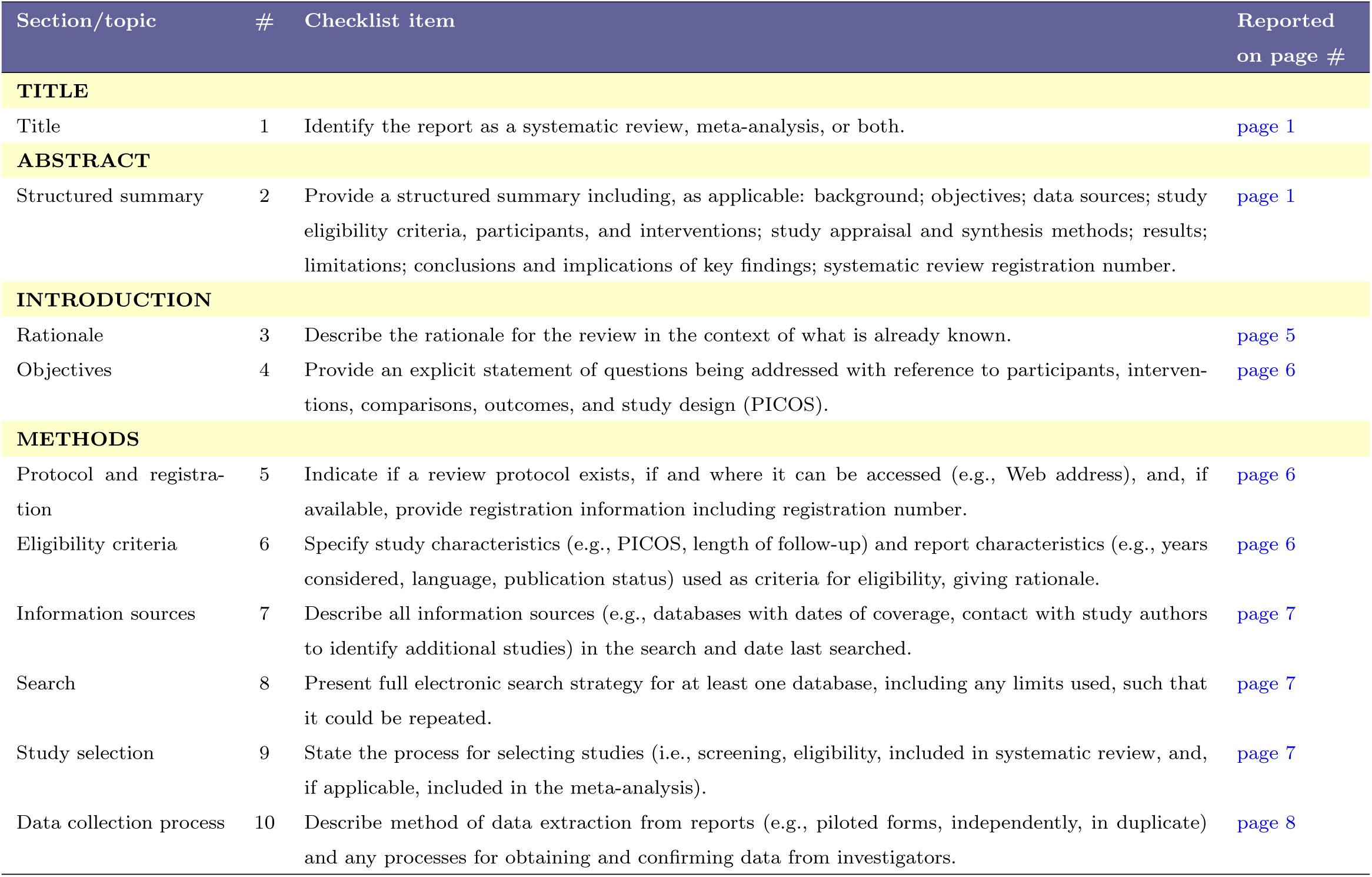
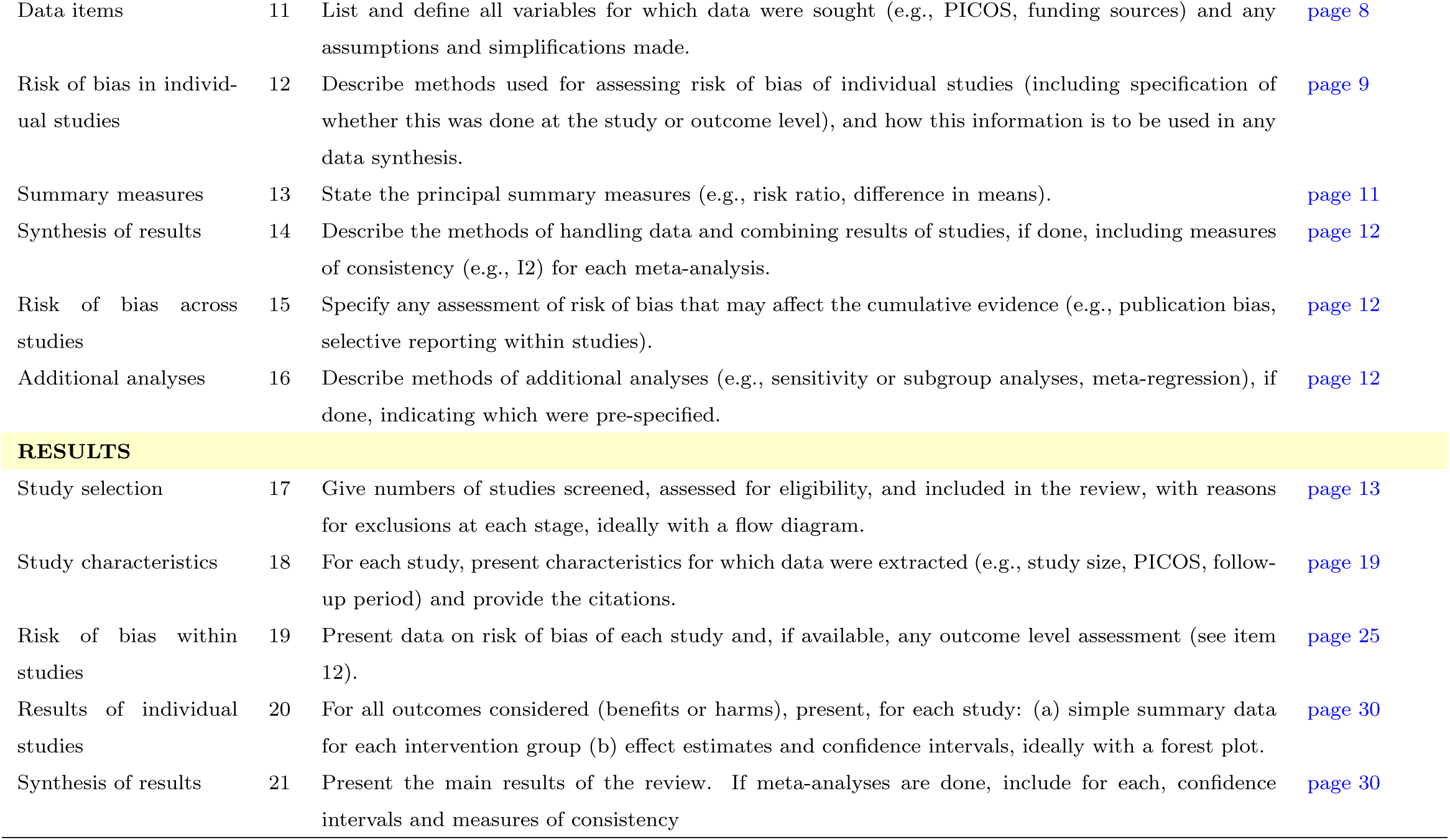
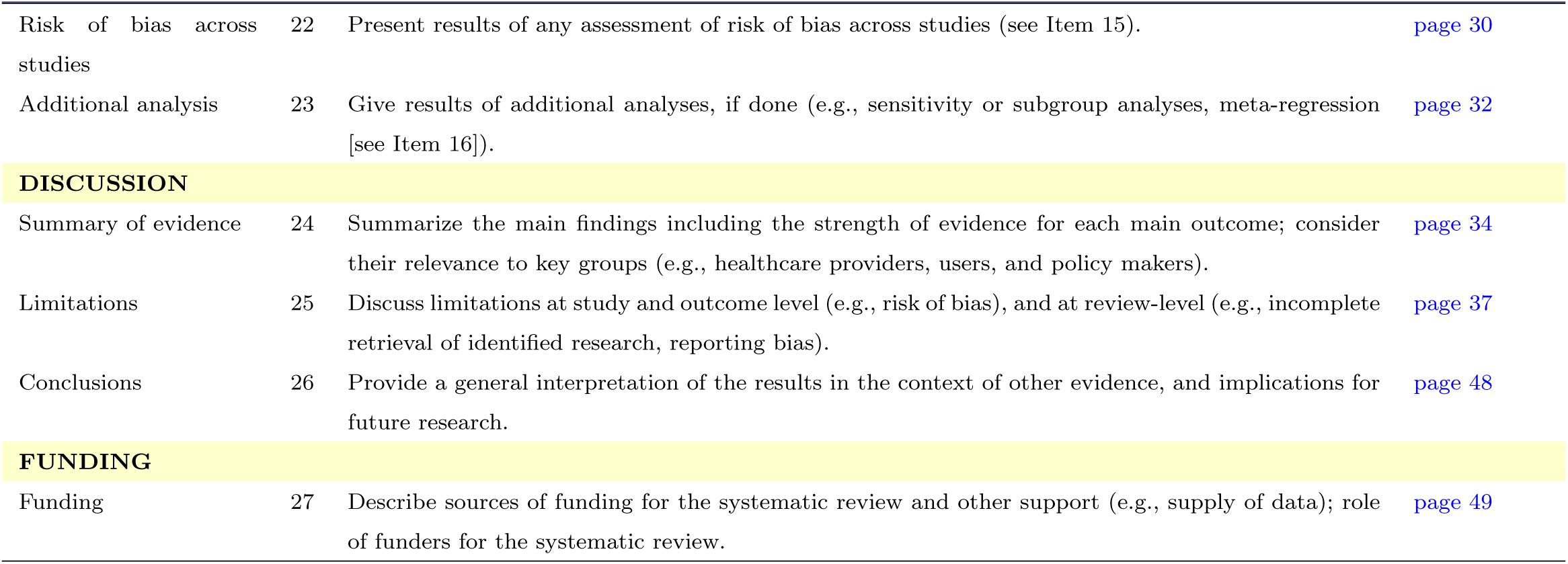
From: Moher et al. (2009)

## Appendix C. Prediction performance metrics

Given a set of true labels ***y***, and a set of predictions 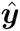, several performance metrics can be defined.

### Appendix C.1. Continuous valued labels

The Pearson correlation coefficient, defined as

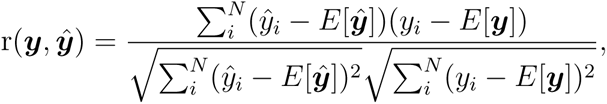

is the most popular performance metric for regression of continuous valued labels.

Other metrics include the MSE,

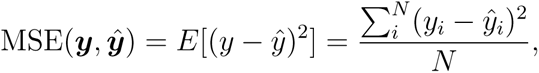

the MAE,

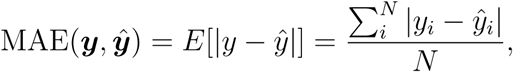

and Spearman rank correlation coefficient,

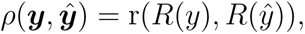

defined in terms of Pearson’s, but based on ranks instead of values, as denoted by the rank function *R*(*·*).

A few more metrics are linked to the MSE. These include the coefficient of determination, or squared deviance, R-squared,

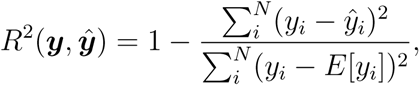

the RMSE,

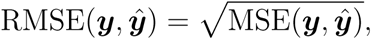

the NRMSE,

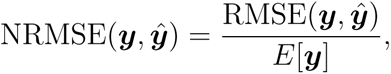

and the nRMSD,

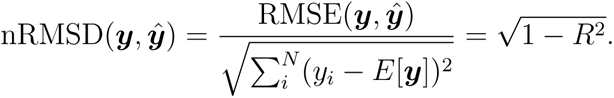

### Appendix C.2. Binary valued labels

In our sample, the only reported performance metric for binary valued labels *y_i_ ∈ {*0, 1*}* was the AUC, defined mathematically as

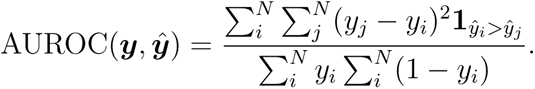

Notice that (*y_j_ − y_i_*)^2^ *≡* 1 only when *y_i_ ∈* = *y_j_*, being 0 otherwise.

## Appendix D. Adjusted TRIPOD checklist

**Table D.5:** Adjusted TRIPOD checklist for reporting quality assessment

| Section/Topic | Item | Checklist Item | Page |
| --- | --- | --- | --- |
| Title and abstract |  |  |  |
| Title | 1 | Identify the study as developing and/or validating a multivariable prediction model, the target population, and the outcome to be predicted. |  |
| Abstract | 2 | Provide a summary of objectives, study design, setting, participants, sample size, predictors, outcome, statistical analysis, results, and conclusions. |  |
| Introduction |  |  |  |
| Background and objectives | 3a | Explain the medical context (including whether diagnostic or prognostic) and rationale for developing or validating the multivariable prediction model, including references to existing models. |  |
|  | 3b | Specify the objectives, including whether the study describes the development or validation of the model or both. |  |
| Methods |  |  |  |

**Table D.5 continued from previous page**
| Section/Topic | Item | Checklist Item | Page |
| --- | --- | --- | --- |
| Source of data | 4a | Describe the study design or source of data (e.g., randomized trial, cohort, or registry data), separately for the development and validation data sets, if applicable. |  |
|  | 4b | Specify the key study dates, including start of accrual; end of accrual; and, if applicable, end of follow-up. |  |
| Participants | 5a | Specify key elements of the study setting (e.g., primary care, secondary care, general population) including number and location of centres. |  |
|  | 5b | Describe eligibility criteria for participants. |  |
|  | 5c | Give details of treatments received, if relevant. |  |
| Outcome | 6a | Clearly define the outcome that is predicted by the prediction model, including how and when assessed. |  |
|  | 6b | Report any actions to blind assessment of the outcome to be predicted. |  |

**Table D.5 continued from previous page**
| Section/Topic | Item | Checklist Item | Page |
| --- | --- | --- | --- |
| Predictors | Adjusted<br>7a | Clearly define all predictors used in developing or validating the ML model, including how and when they were measured. |  |
|  | 7b | Report any actions to blind assessment of predictors for the outcome and other predictors. |  |
| Sample size | 8 | Explain how the study size was arrived at. |  |
| Missing data | 9 | Describe how missing data were handled (e.g., complete-case analysis, single imputation, multiple imputation) with details of any imputation method. |  |
| Statistical<br>analysis<br>methods | 10a | Describe how predictors were handled in the analyses. |  |
|  | Adjusted<br>10b | Specify type of model, all model-building procedures (including any predictor selection, hyperparameter selection if needed), and method for internal validation. |  |
|  | 10d | Specify all measures used to assess model performance and, if relevant, to compare multiple models. |  |

**Table D.5 continued from previous page**
| Section/Topic | Item | Checklist Item | Page |
| --- | --- | --- | --- |
| Risk groups | 11 | Provide details on how risk groups were created, if done. |  |
| Results |  |  |  |
| Participants | 13a | Describe the flow of participants through the study, including the number of participants with and without the outcome and, if applicable, a summary of the follow-up time. A diagram may be helpful. |  |
|  | 13b | Describe the characteristics of the participants (basic demographics, clinical features, available predictors), including the number of participants with missing data for predictors and outcome. |  |
| Model development | 14a | Specify the number of participants and outcome events in each analysis. |  |
|  | 14b | If done, report the unadjusted association between each candidate predictor and outcome. |  |

**Table D.5 continued from previous page**
| Section/Topic | Item | Checklist Item | Page |
| --- | --- | --- | --- |
| Model specification | Adjusted 15a | Present the full prediction model to allow predictions for individuals (i.e. links to the final model online (coding of predictors, code and final parameters/coefficients, and with the architecture described in full in the article)). |  |
|  | 15b | Explain how to use the prediction model. |  |
| Model performance | 16 | Report performance measures (with CIs) for the prediction model. |  |
| Discussion |  |  |  |
| Limitations | 18 | Discuss any limitations of the study (such as nonrepresentative sample, few events per predictor, missing data). |  |
| Interpretation | 19b | Give an overall interpretation of the results, considering objectives, limitations, and results from similar studies, and other relevant evidence. |  |
| Implications | 20 | Discuss the potential clinical use of the model and implications for future research. |  |
| Other information |  |  |  |

Table D.5: Adjusted TRIPOD checklist for reporting quality assessment
| Section/Topic | Item | Checklist Item | Page |
| --- | --- | --- | --- |
| Supplementary information | 21 | Provide information about the availability of supplementary resources, such as study protocol, Web calculator, and data sets. |  |
| Funding | 22 | Give the source of funding and the role of the funders for the present study. |  |

## Appendix E. Synthesis of meta-analytic results

**Figure E.8:**
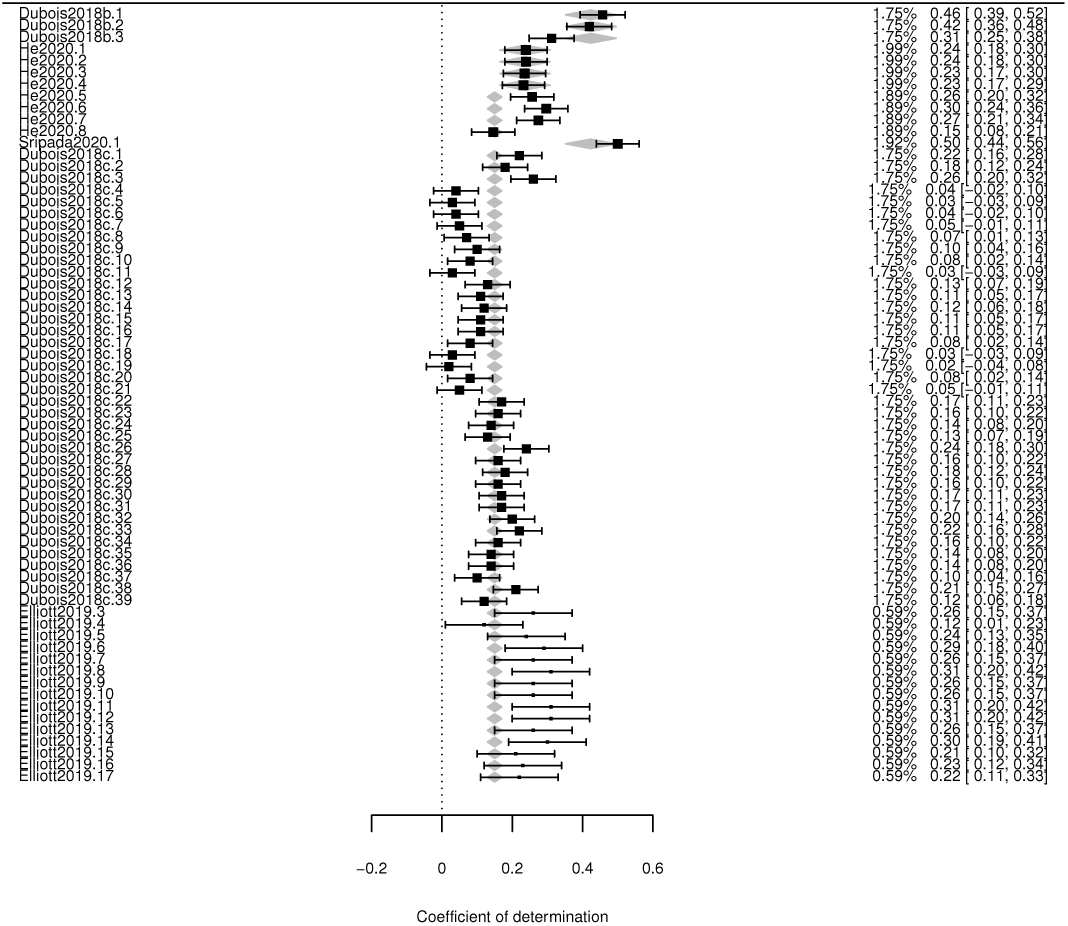
Forest plot for the correlation coefficient meta-analysis.

**Figure E.9:**
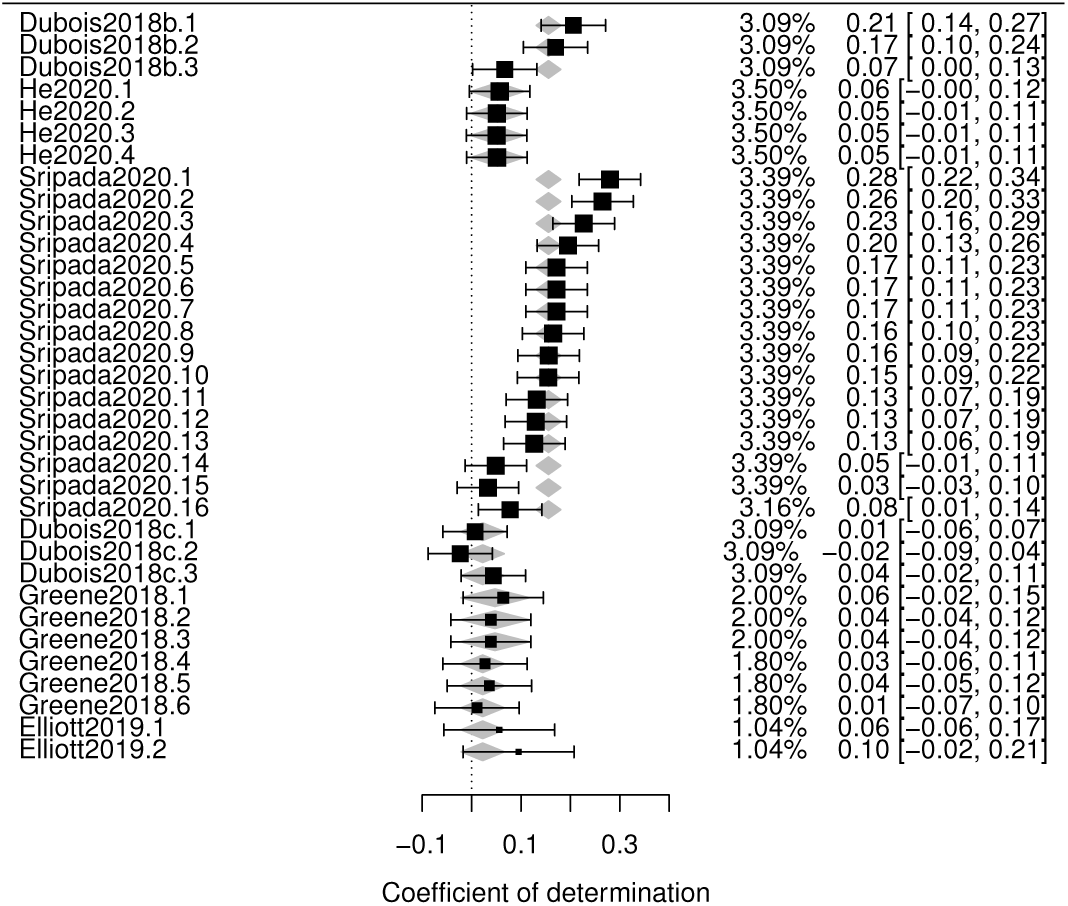
Forest plot for the R-squared meta-analysis.

## Appendix F. Risk of bias across studies

**Figure F.10:**
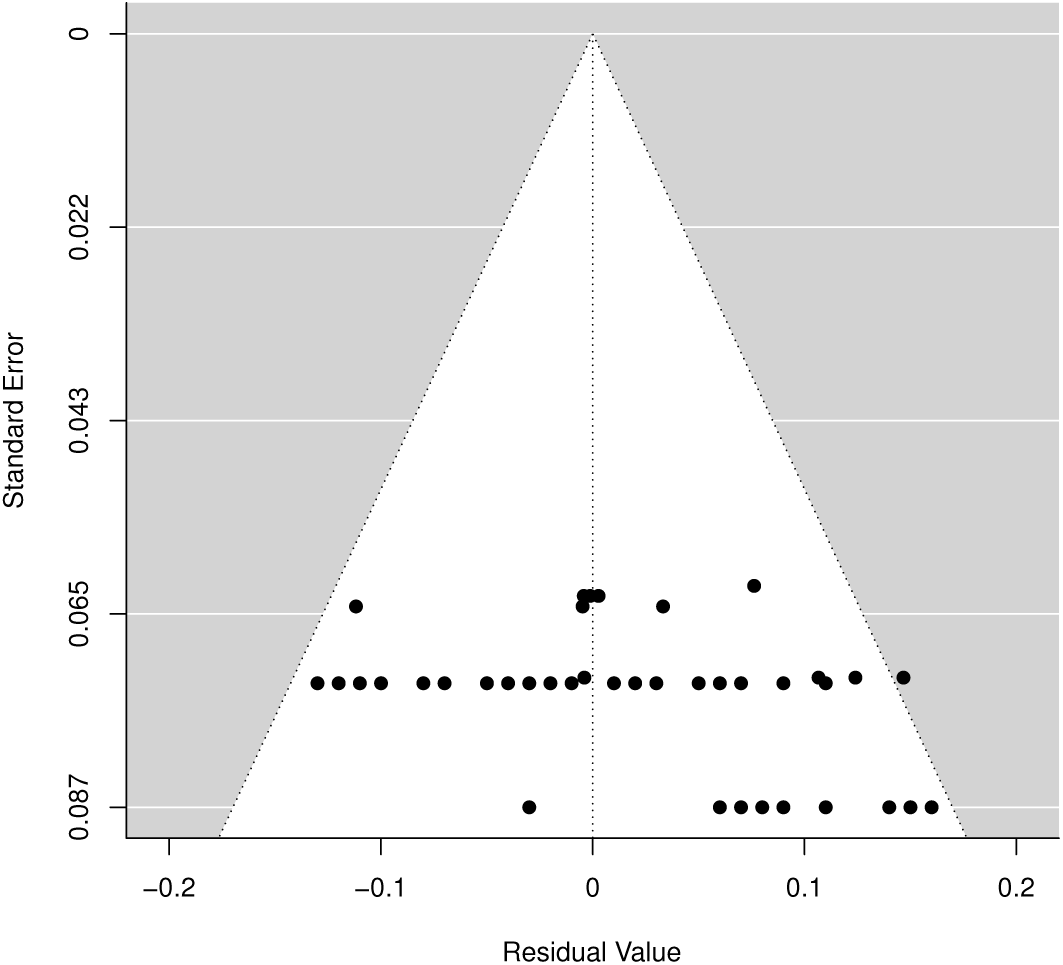
Funnel plot for the correlation coefficient meta-analysis.

**Figure F.11:**
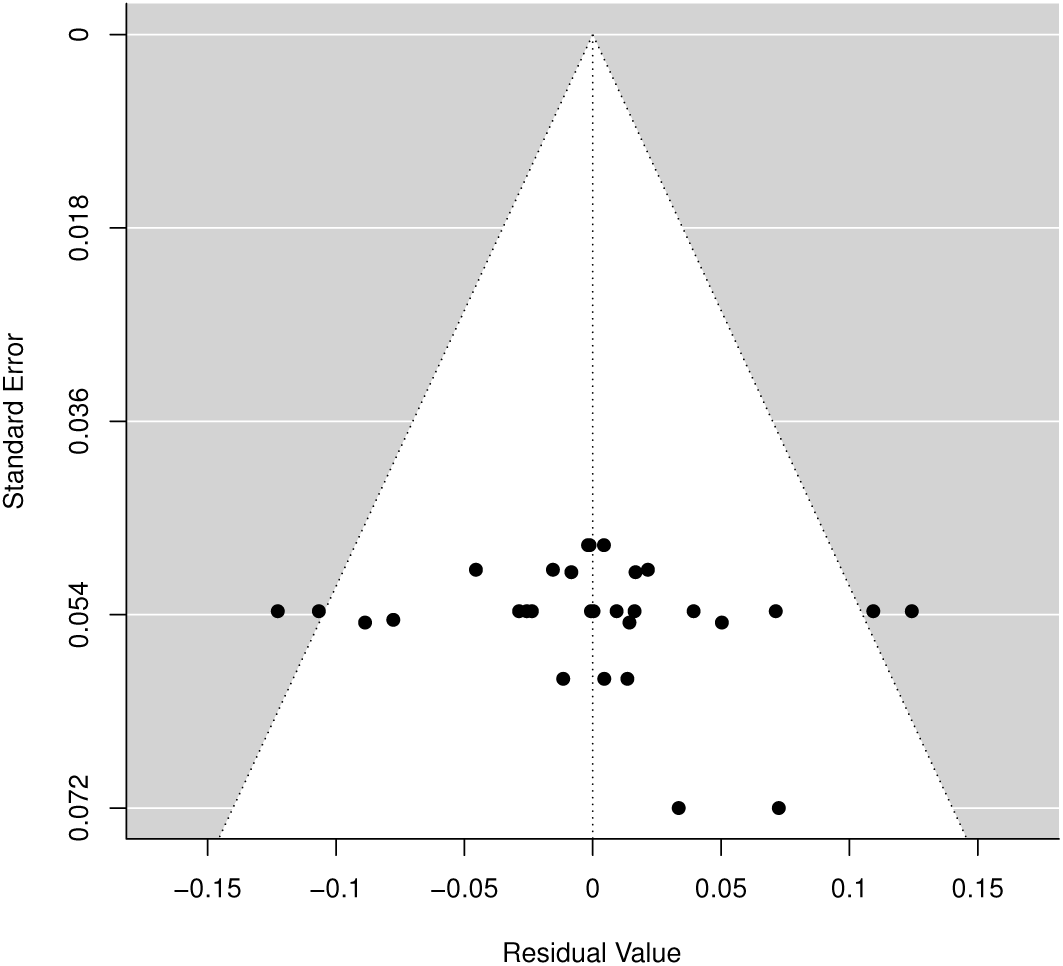
Funnel plot for the R-squared meta-analysis.

## Notes

https://osf.io/5qe64/

### Competing Interest Statement

The authors have declared no competing interest.

